# The influence of nonlinear resonance on human cortical oscillations

**DOI:** 10.1101/2025.06.27.661950

**Authors:** Ying Wang, Min Li, Ronaldo García Reyes, Maria L. Bringas-Vega, Ludovico Minati, Michael Breakspear, Pedro A. Valdes-Sosa

## Abstract

A longstanding debate in neuroscience concerns whether macroscale brain signals can be described as purely linear Gaussian processes or they harbor the more complex statistics of nonlinear dynamics. We introduce BiSpectral EEG Component Analysis (BiSCA), a framework unifying power spectral and bispectral analysis to test for nonlinearity by identifying inter-frequency harmonic relationships and disambiguating them from superimposed components. In particular, simulations confirm that it is able to separate valid nonlinearity from non-Gaussianity, which is a common confound. Applying this test to two large human brain recordings datasets (1,771 intracranial channels and 960 individuals’ scalp EEG), we find the brain’s broadband, aperiodic background behaves as a linear, Gaussian process, while narrowband Rho oscillatory --including the Alpha and Mu rhythms --are the primary source of cortical nonlinearity, exhibiting significant quadratic cross-frequency coupling. Both recordings show significant departures from linear Gaussian behavior. We observe a clear dissociation between signal power and nonlinearity. While the occipital Alpha rhythm dominates the power spectrum, the strongest nonlinear signatures arise from the less dominant parietal Mu rhythm. These findings suggest that nonlinear resonance is pervasive in cortical signals, primarily expressed through resonant oscillations rather than aperiodic activity. Without accounting for these intrinsic nonlinear interactions, a principled understanding of neural activity will be incomplete.

## 1. Introduction

Can mesoscale brain dynamics, as reflected in resting state EEG (rsEEG) and fMRI (rsfMRI), be described entirely as linear, Gaussian stochastic processes? Recent studies, such as Nozari et al.(2024) ^[1]^ and Raffaelli et al.(2024) ^[2]^, argue that linear time-series models suffice to describe these signals. The authors of these studies suggest that the intricate nonlinearities observed at the microscale—by neurons, synapses, and neural masses—are effectively “averaged out” at larger scales. If this observation were true, the physics of the observation process would imply that mesoscale signals lack observable nonlinear signatures, limiting our ability to infer microscale dynamics from macroscopic measurements. Conversely, empirical refutation of this claim would imply that neuronal signals possess correlations at all scales, sufficient to avoid the diffusive “washing out” of macroscopic averaging and thus retain nonlinear properties at large scales ^[3]^.

The high temporal resolution of the scalp or intracranial resting-state EEG (rsEEG) (Fig. 1A) makes it particularly suitable for investigating dynamic neural processes. Therefore, reports asserting linearity or Gaussianity are somewhat surprising since substantial evidence indicates that nonlinear dynamics are observable in rsEEG signals ^[4,5]^. Nonlinear dynamics in neural systems, as evidenced by phenomena in the frequency domain such as nonlinear resonance, cross-frequency coupling (CFC), and phase-amplitude coupling (PAC), are critical for understanding the complex oscillatory interactions observed in resting-state EEG (rsEEG) and task-related EEG signals. One of these common features of nonlinear neural oscillations is their non-sinusoidal waveform shape, which is abundant in human electrophysiological recordings and reflects the complex biophysical dynamics of underlying neural generators ^[6]^, as modeled by early neural mass models ^[7–10]^. Conventional spectral analysis ignores non-sinusoidal structure. These carry important physiological information, and can distinguish between different behavioral states ^[6,11–13]^. Such complex waveforms inevitably generate harmonic components in the frequency spectrum ^[14–18,18–22]^. This asymmetric nonlinearity or quadratic phase coupling can be described by bicoherence ^[7,23]^. This emphasis on Bicoherence is bolstered by recent theoretical and empirical work that shows that other popular measures, such as phase-amplitude coupling, are actually equivalent to it. This measure thus provides a unified framework for understanding nonlinear resonance in neural systems^[24]^. Furthermore, large-scale brain network models have shown that cross-frequency coupling emerges naturally from the nonlinear dynamics of interconnected neural populations, supporting the view that nonlinear resonance is a fundamental organizing principle of brain activity ^[25]^. Critically, if macroscopic signals were limited to low-order statistical properties (e.g., mean, variance), none of these nonlinear resonance phenomena would have been observed. Moreover, widely and successfully used techniques like Independent Component Analysis (ICA) — which relies on higher-order moments—would fail to separate signal sources.

**Fig. 1.**
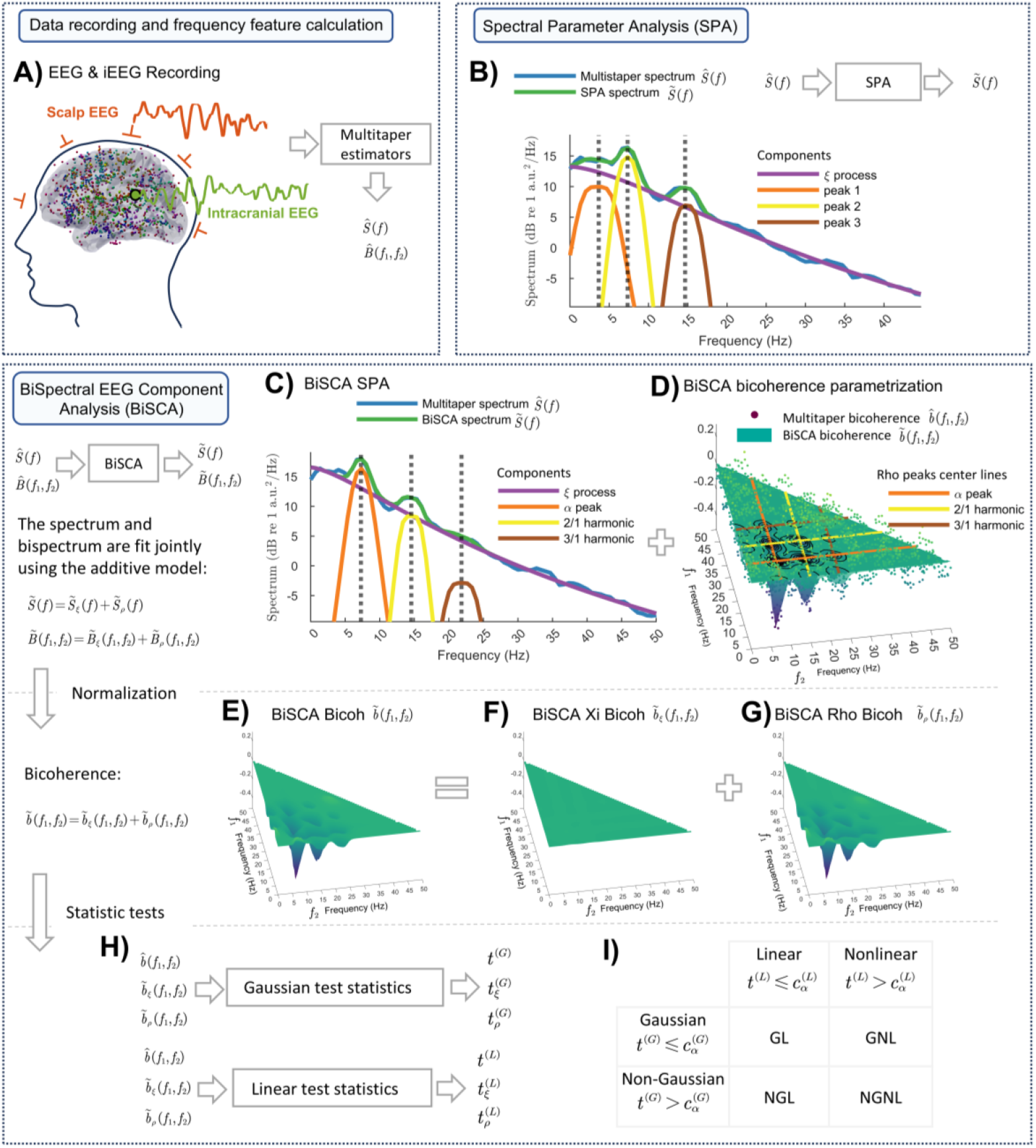
Overview of the BiSpectral Components Analysis (BiSCA) framework. (A) Data recording (iEEG and scalp EEG) and calculation of Multitaper spectral and bispectral estimators. (B) Spectral Parameter Analysis (SPA) decomposing the power spectrum into individual components. (C) BiSCA model fitting the power spectrum, capturing harmonic relationships. (D) BiSCA model fitting the bicoherence to parametrize nonlinear interactions. (E-G) Additive decomposition of the BiSCA bicoherence into (E) total bicoherence, (F) the Xi-component, and (G) the Rho-component. (H) Derivation of statistics for Gaussianity and linearity tests from bicoherence measures. (I) Table for classifying a signal as Gaussian/Non-Gaussian and Linear/Nonlinear based on statistical thresholds.

The debate upon the linearity or nonlinearity of the EEG tends to be muddied by the fact that a critical concept is ignored, namely, the distinction between linearity, that is, the nature of a system’s temporal evolution, and Gaussianity, that is, the statistical distribution of its variables. Even when a system is linear, for example, an autoregressive model, when driven by non-Gaussian inputs (e.g., skewed or heavy-tailed noise), it can yield non-Gaussian outputs. On the other hand, a nonlinear system can also produce approximately Gaussian output. This can happen due to the summation of multiple independent processes, where the Central Limit Theorem scenario applies. Non-Gaussianity can also stem from the intrinsic dynamics of the system ^[26]^. In fact, nonlinear systems frequently exhibit pronounced non-Gaussianity, such as multimodal or skewed distributions. This occurs especially in the presence of harmonic coupling or multiplicative feedback. Consequently, neural dynamics can embody any combination of linearity/nonlinearity and Gaussianity/non-Gaussianity. Simple inspection of a power spectrum of these combinations of generative mechanisms, as demonstrated in Supplementary Materials S.3. Distinguishing the generative mechanisms requires moving beyond simple distributional analysis or the tests upon the spectrum and directly testing for nonlinear interactions. Prime examples of nonlinear behavior to search for are nonlinear resonance phenomena like phase coupling and harmonic generation.

It is also important to recognize that the EEG signal is likely a mixture of different types of processes and that Gaussianity or linearity (or their absence) may vary across these processes. By dissecting the contributions of various subprocesses—such as oscillatory rhythms, aperiodic activity, and noise—it becomes possible to identify which components adhere to linear dynamics and Gaussian statistics and which deviate due to nonlinear interactions or non-Gaussian inputs.

A successful area of research for decomposing the EEG into such processes is based on the EEG second-order cumulant *S* (*f*) or spectrum for frequency *f*. For strictly stationary processes without long-term memory ^[27]^, the distribution of optimized estimators *Ŝ*(*f*) (for example, with Multitaper) completely characterizes linear time-invariant signals driven by noise. The separation of an EEG *Ŝ*(*f*) into distinct components defined by simple functional forms, each determined by only a few parameters, is known as **Spectral Parameter Analysis** (**SPA**) (Fig. 1B & Supplementary S.2). An early example of SPA is Pascual-Marqui et al. (1987)^[28]^, who proposed the Xi-Alpha (ξα) model. They showed that EEG spectra have two dominant components: an aperiodic Xi (ξ) component and multiple peaks characterizing rhythmic activity (which we will call generically Rho components or ρ), the best known being the Alpha (α) component (the Peak 1 in Fig. 1B). The FOOOF(Fitting Oscillations & One Over F) model is a recent example of SPA ^[29]^. Not only do these components have distinct behavioral correlates ^[30–32]^, they may also derive from distinctive physiological processes ^[33,34]^.

However, SPA, which relies on second-order moments (the power spectrum), only provides a sufficient characterization for linear Gaussian systems. For such systems, all higher-order cumulants are zero. Consequently, SPA is thus blind to nonlinear dynamics. For this type of activity, higher-order spectral tools, such as the bispectrum *B* (*f*_1_*f*_2_) and its normalized version, the bicoherence *b* (*f*_1_*f*_2_), between frequencies *f*_1_ and *f*_2_, are required to detect phenomena such as harmonic generation, phase coupling, or other nonlinear interactions. Analysis of nonlinear time series (multiple lags) is therefore needed. Another fundamental problem of state-of-the-art SPA is that the harmonic relationships between second-order spectral peaks are not explicitly modeled despite their being ubiquitous in EEG recordings.

We introduce **BiSCA** (**BiS**pectral EEG **C**omponent **A**nalysis), a generalized model of EEG spectral components designed to detect and quantify nonlinear resonance. It thus addresses the limitations of the current practice of relying solely on isolated second-order spectral peaks. Instead, BiSCA provides a combined model specifically designed to identify nonlinear resonance phenomena in neural signals, also modeling harmonically related second-order peaks and their corresponding bispectral counterparts. Furthermore, BiSCA parameter estimation is based on maximum likelihood, enabling direct statistical testing of linearity and Gaussianity for each component (Fig. 1E-G). Following in-silico validation, we demonstrate how the BiSCA model reveals substantial nonlinearities and non-Gaussian characteristics in EEG data using 1,771 resting-state intracranial EEG (iEEG) recordings from 106 patients.

## 2. Results

### 2.1 Overview of BiSpectral Components Analysis (BiSCA)

The BiSpectral Components Analysis (BiSCA) framework is designed to uncover nonlinearity and non-Gaussianity in resting-state EEG signals. Fig. 1 provides a schematic overview of this methodology. The analysis pipeline begins with electrophysiological data, including intracranial (iEEG) and scalp EEG recordings (Fig. 1A). From these timeseries, we compute initial data summaries using Multitaper estimators to obtain the power spectrum, denoted as *Ŝ*(*f*), and the bispectrum, 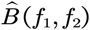.

A conventional approach, Spectral Parameter Analysis (SPA), decomposes the power spectrum 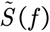 on either a natural or logarithmic scale into a linear superposition of an aperiodic background component and several oscillatory peaks (Fig. 1B). However, SPA does not account for higher-order statistics, a limitation that prevents it from modeling the harmonic relationships between these peaks.

The BiSCA model extends this approach by jointly fitting both the power spectrum and the bispectrum (Fig. 1C&D). This joint modeling explicitly parameterizes the nonlinear resonances that manifest as harmonic relationships in the spectrum and corresponding peaks in the bispectrum. The model additively separates the signal into components, such as the aperiodic Xi (ξ) process and the oscillatory Rho (ρ) process. Fig. 1C shows the BiSCA model fits to the power spectrum, identifying a fundamental alpha peak and its harmonics. Fig. 1D illustrates the corresponding fit to the bicoherence 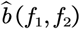 (the normalized bispectrum 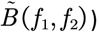, where harmonic relationships are visible as distinct patterns. A key feature of BiSCA is the ability to decompose the total bicoherence into contributions from individual components (Fig. 1E-G). For instance, the total bicoherence (Fig. 1E) can be separated into the bicoherence arising from the Xi-component (Fig. 1F) and the Rho-component (Fig. 1G), isolating the source of nonlinearity.

Finally, to formally test for the presence of such phase-coupled interactions, statistical tests are applied based on the bicoherence estimates to assess Gaussianity and linearity (Fig. 1H). These tests are conducted jointly, are sensitive to quadratic nonlinearity, and distinguish whether deviations from a Linear Gaussian process stem from non-Gaussian inputs or from the system’s nonlinear dynamics. As summarized in Fig. 1I, this parallel assessment classifies a signal as Gaussian Linear (GL), Non-Gaussian Linear (NGL), Gaussian Nonlinear (GNL), or Non-Gaussian Nonlinear (NGNL), the latter three categories (NGL, GNL, and NGNL) all manifest as non-Gaussian signals at the observational level.

### 2.2 Simulation Illustrating Asymmetric Waveforms and Quadratic Nonlinearity

To build a clear intuition for the bicoherence-based tests central to the BiSCA framework, we use a targeted simulation to illustrate what they specifically detect: quadratic (or even-order) nonlinearities, which manifest as waveform asymmetry. This comparison highlights the specific features our test detects, rather than merely differentiating a linear-Gaussian process from a nonlinear one.

The analysis begins with an example of the “wicket waves” ^[35]^ from a 2.5-second iEEG segment Fig. 2A, waveforms known for their strong asymmetry. A Nonparametric Nonlinear Autoregressive (NNAR) model was fitted to this data and pruned to define its core dynamics. This model was then used to generate two types of long-running simulations.

**Fig. 2.**
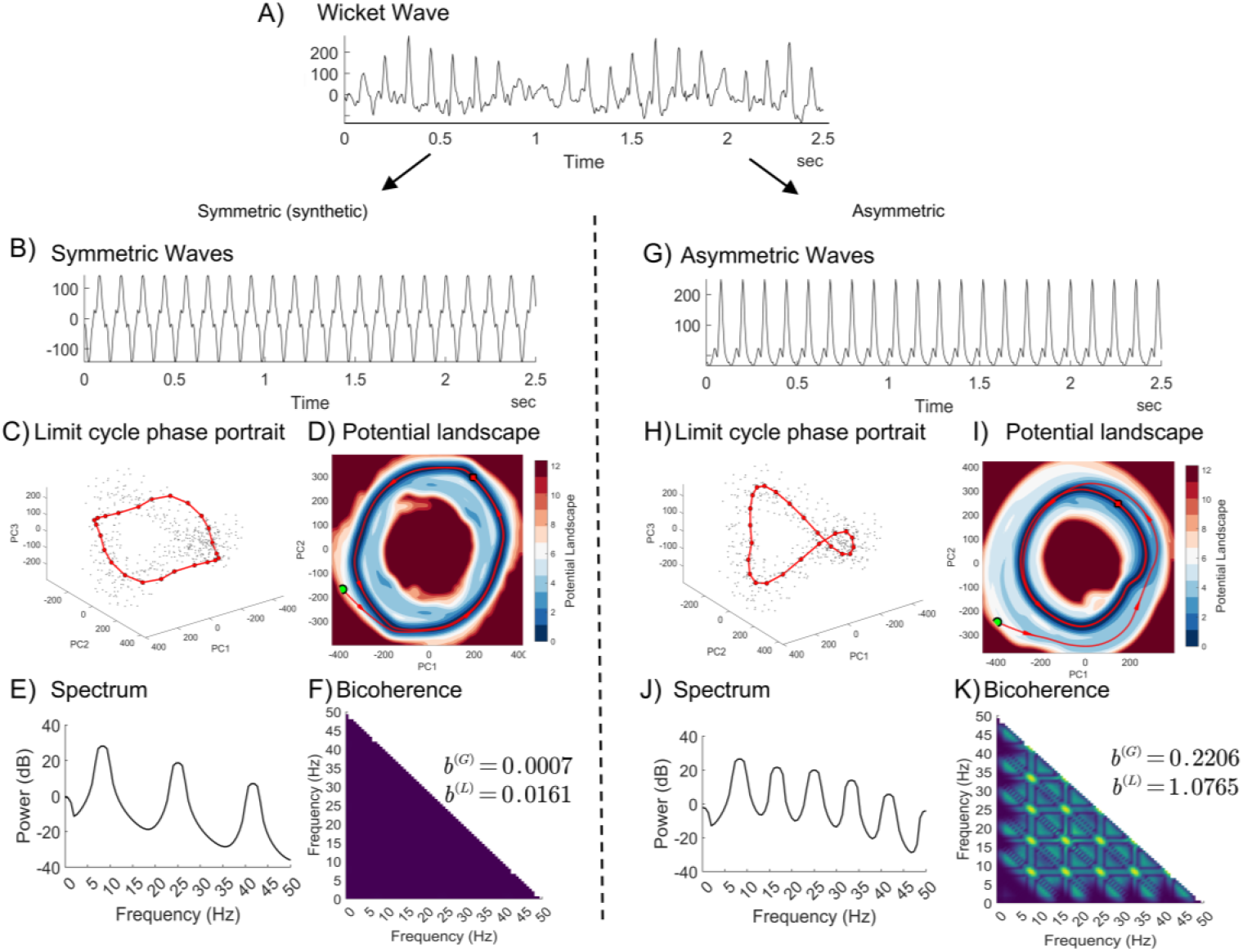
Comparison of asymmetric and symmetric nonlinear systems to illustrate quadratic nonlinearity. The simulations are derived from an NNAR model fitted to an iEEG Wicket Wave segment (A). Left Panel (B-F): Synthetic symmetric system. (B) Symmetric waveform; (C) symmetric 3D limit cycle phase portrait (red line) with pruned model states (gray dots) used for the model fit; (D) symmetric 2D potential landscape; (E) corresponding power spectrum exhibiting only odd-order harmonics; and (F) negligible bicoherence. Right Panel (G-K): Asymmetric system. (G) Asymmetric waveform; (H) asymmetric 3D limit cycle; (I) asymmetric 2D potential well; (J) power spectrum containing both even and odd harmonics; and (K) significant bicoherence peaks indicating quadratic phase coupling.

The right panel of Fig. 2 (G-K) shows a simulation preserving the original asymmetry of the wicket waves. For this system, the estimated 2D potential landscape of the dynamics reveals a distinctly asymmetric basin of attraction (Fig. 2I). The shape of this potential well geometrically constrains the system’s trajectory thus forcing the system into the distorted, non-uniform limit cycle shown in the 3D phase portrait (Fig. 2H). This dynamical structure generates the asymmetric morphology of the time-domain waveform (Fig. 2G), reflected in the spectrum (Fig. 2J). Note that the spectrum contains both even and odd harmonics of the fundamental frequency (around 8.7 Hz). This harmonic structure is reflected by quadratic phase coupling among the harmonics, signaled by as high-valued peaks in the bicoherence plot Fig. 2K, quantified by a bicoherence magnitude of *b*^(*L*)^ = 1.0765.

Conversely, the left panel (Fig. 2B-F) shows a synthetic simulation where the waveform is artificially symmetrized by enforcing odd symmetry in the model dynamics (*F* (− ***x***) = − *F* (***x***), see details in supplementary). This process generates a perfectly symmetric, albeit still non-sinusoidal, waveform (Fig. 2B). The corresponding state-space representations are also symmetric. The geometry of the dynamics corresponds to a concentric 2D potential well (Fig. 2D) forcing the trajectory to trace a symmetric 3D limit cycle (Fig. 2C), which is plotted along with the pruned states (coreset, gray dots) used by the model. Critically, the spectrum of this symmetric system (Fig. 2E) contains only odd-order harmonics. Since bicoherence quantifies quadratic (even-order) interactions, the absence of even harmonics results in a statistically negligible bicoherence value (Fig. 2F), with a magnitude of *b*^(*L*)^ = 0.0161. This directly addresses the observation that a non-sinusoidal waveform can have minimal bicoherence if its nonlinearity is purely symmetric (odd-order). This lack of sensitivity to even harmonic coupling will not affect our conclusions since analyzing higher order statistics would be more cumbersome and only increase the rejection of linear behavior.

These two simulations illustrate how the geometric asymmetry in the state-space representations of the estimated potential landscape and corresponding limit cycle show up in the bifrequency plane as quadratic phase coupling quantifiable by bicoherence. Such dynamic behavior is inaccessible to a linear system, which is fundamentally constrained to a symmetric, parabolic potential well, producing only an elliptical trajectory and exhibiting negligible bicoherence. This example illustrates how our test effectively identifies quadratic nonlinearities and distinguishes them from other forms of nonlinear dynamics.

### 2.3 Nonlinear and non-Gaussian cortical dynamics in EEG/iEEG revealed by Bicoherence

To assess the prevalence of nonlinear and non-Gaussian dynamics in the EEG and iEEG datasets, we applied statistical tests for linearity and Gaussianity. These tests are based on Multitaper bicoherence estimates 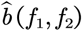, employing a modified Hinich test ^[36]^The thresholds for the tests were adjusted for multiple comparisons using the Benjamini-Hochberg FDR procedure (q=0.001). Each recording was classified by the joint outcome of two parallel tests—one for linearity and one for Gaussianity. These two results allow us to classify each recording into one of four categories: Gaussian Linear (GL), Non-Gaussian Linear (NGL), Gaussian Nonlinear (GNL), or Non-Gaussian Nonlinear (NGNL).

The results, summarized in Table 1, indicate widespread deviations of the data from linear Gaussian behavior. A minority of recordings were classified as GL: only 14.7% for EEG and 25.7% for iEEG. Conversely, a substantial majority—85.2% (81.6%+3.6%) of EEG and 74.3% (67.9%+6.4%) of iEEG recordings, respectively—were identified as either nonlinear or non-Gaussian. signals that were strictly linear but non-Gaussian (NGL) were uncommon, accounting for only 3.6% of EEG and 6.4% of iEEG recordings.

**Table 1.** Percentages of EEG and iEEG recordings that are Linear, Nonlinear, Gaussian or non-Gaussian.

|  | EEG |  |  | iEEG |  |  |
| --- | --- | --- | --- | --- | --- | --- |
|  | Linear | Nonlinear | Total | Linear | Nonlinear | Total |
| Gaussian | 14.7 | 39.3 | 54 | 25.7 | 17.1 | 42.7 |
| Non-Gaussian | 3.6 | 42.3 | 46 | 6.4 | 50.9 | 57.3 |
| Total | 18.4 | 81.6 | 100 | 32.1 | 67.9 | 100 |

The spatial distribution of these classifications across electrodes are revealed by topographic maps. These maps exhibit distinctive patterns (Fig. 3A&C). For EEG, linear signals (both Gaussian and non-Gaussian) are most frequently observed at central electrodes (Fig. 3A). In contrast, iEEG shows a predominance of GL signals in the frontal region, while NGL signals are more frequent in the occipital area. GNL activity is strongly expressed in the frontal areas for EEG but is less apparent in iEEG. Nonlinear Non-Gaussian (NGNL) dynamics in EEG show an anterior-to-posterior gradient, peaking at occipital electrodes, whereas in iEEG, they are concentrated centrally with a right-hemispheric asymmetry in the occipital lobes.

**Fig. 3.**
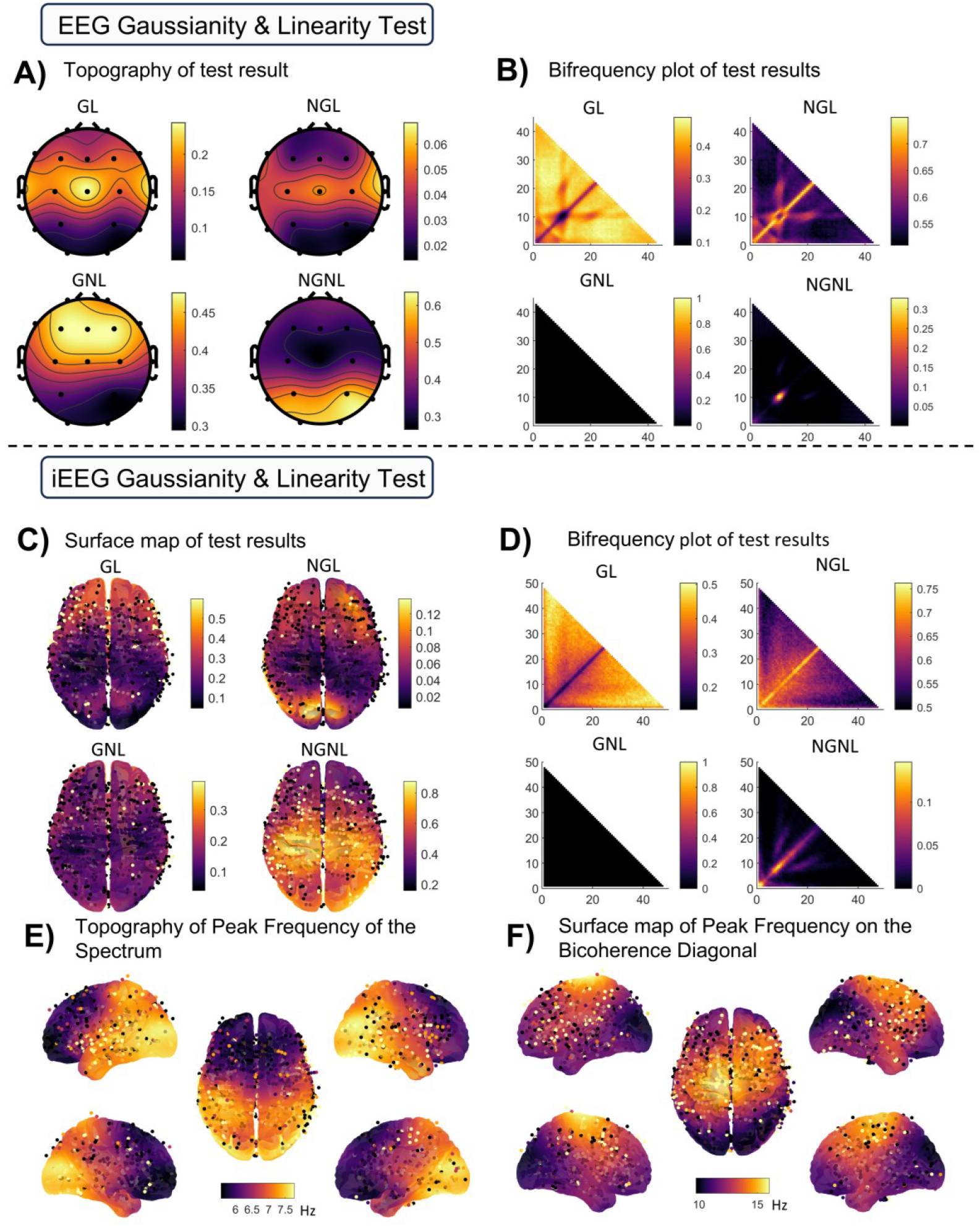
Results of Gaussianity and linearity tests applied to iEEG and scalp EEG recordings. (A) The topography map of the proportion of test classes of EEG data; (B) the proportion of test class of EEG along the bifrequency panel; (C) the surface plot interpolated by the statistical test result (0: accept, 1: reject) of iEEG channels; (D) the proportion of test class of iEEG along the bifrequency panel; (E-F) the spatial distribution of the frequency for the maximum spectrum peak and bicoherence peak after removing the Xi trend.

An alternative perspective is offered by plotting the proportion of test result types over the triangular bifrequency domain (*f*_1_*f*_2_) and *f*_1_ + *f*_2_ ⩽ 50Hz (corresponding to the bicoherence principal domain, here after bifrequency panel) (Fig. 3B&D). Note that here, though we deal with statistics for each (*f*_1_*f*_2_) pair, the threshold for deciding tests is for the family of tests over the entire triangular plane, corrected with FDR. Also, by construction, the Gaussianity threshold 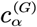 and the 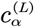 linearity threshold are related as 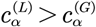, for individual tests, resulting in a zero proportion of nonlinear but Gaussian results (lower left of Fig. 3B&D). A diagonal ridge of Non-Gaussian Linear test results suggests time domain asymmetries of waveforms for all frequencies. Note, however, the off-diagonal bands of interference coupling for the Alpha frequency harmonics suggest frequency doubling and intermodulation effects. Note the NGNL plot for EEG reveals a localized region of high proportion near (10 Hz, 10 Hz) associated with Nonlinearity and Non-Gaussianity. This result indicates evidence of intra-band nonlinear coupling in the alpha rhythm. The bifrequency distribution for iEEG (Fig. 3D) shows a qualitatively similar structure to the EEG results. However, these patterns in the iEEG data are more diffusely distributed over the frequency triangle compared to scalp EEG data.

It is instructive to view the topographies of the peak frequencies at which the spectrum is maximum, 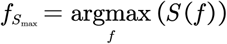, and the frequency at which the bicoherence is maximal, determined using the maximum of the main diagonal bicoherence 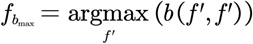, indicates that where the frequency-doubling occurs (Fig. 3E&F). The latter is restricted to the bispectral values on the main diagonal of the principal domain. The occipital and parietal areas possess these local maxima in the alpha band, while the frontal region is in the theta band. The peak frequency of the bicoherence is higher than the peaks found by the spectrum.

### 2.4 Nonlinear resonances manifest in Rho peaks, while the Xi process remains linear and Gaussian

To understand how each process contributes to the signal’s overall statistical properties, we analyzed them separately. The aperiodic Xi process consistently behaved as a linear, Gaussian process: its Gaussianity statistic 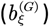 remained well below the significance threshold (Fig. 4A), and its nonlinearity statistic 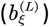 rarely exceeded the threshold (Fig. 4B).

**Fig. 4.**
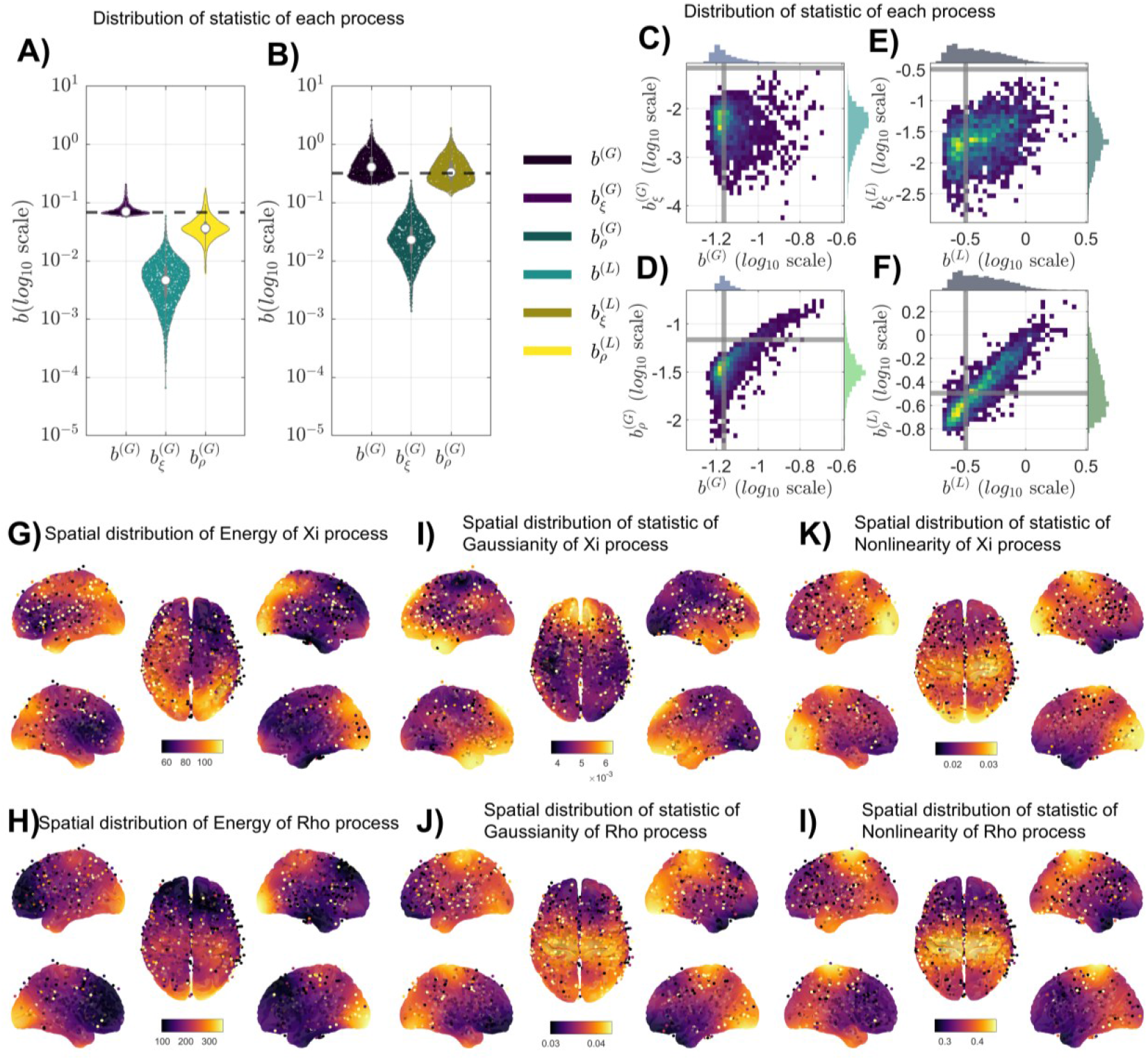
Statistical features of iEEG signals decomposed into Xi trends and periodical Rho components. (A-B) Violin plots showing the distribution of the (A) Gaussianity statistic b^(G)^ and (B) nonlinearity (bicoherence) statistic b^(L)^ for the full signal, the aperiodical Xi process (ξ), and the periodical Rho processes (ρ). The dashed line indicates the threshold for statistical significance. **(C-F)** Scatter plots showing the relationship between the full signal and its components. Plots (C-D) show the correlation between the Gaussianity statistics of the full signal and the (C) Xi processes and the (D) Rho process. Plots (E-F) show the correlation between the nonlinearity statistics of the full signal and the (E) Xi processes and the (F) Rho process. (G-L) Spatial distribution maps showing key statistics for the aperiodic Xi and Rho processes. (G-H) Spatial distribution of energy. (I-J) Spatial distribution of the Gaussianity statistic b^(G)^. (K-L) Spatial distribution of the nonlinearity statistic b^(L)^. Red indicates higher values, while blue indicates lower values.

In stark contrast, the periodic Rho processes were the primary source of nonlinearity. While being largely Gaussian (with 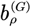 below threshold), their nonlinearity statistic 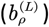 was significantly higher than the threshold, mirroring the behavior of the full signal. These results clearly indicate that the nonlinearity detected in the composite iEEG signal is driven by its rhythmic components, not its aperiodic background.

We further examined how the statistical properties of the full signal are driven by its components. The scatterplots reveal a strong correlation between the nonlinearity of the full signal and that of the Rho processes (Fig. 4F), but no such relationship exists for the Xi process (Fig. 4E). Similarly, the full signal’s deviation from Gaussianity is also tied to the Rho components (Fig. 4C-D). An interesting question is the extent to which tests for Linearity and Gaussianity of the original signal are influenced by those of each spectral component. The scatterplots of the tests for both linearity and Gaussianity between the full signal and its components shed light on this issue. A clear relation exists between the two types of tests for the full signal and the Rho processes (Fig. 4F). This is not the case for the Xi process (Fig. 4E). Therefore, the primary factors contributing to the rejection of Gaussianity and linearity in the iEEG appear to be nonlinear resonances related to the Rho processes.

Another crucial aspect of each spectral component is the spatial distribution of their statistical features across the brain (Fig. 4G-L). The energy (sum of the power spectrum over all frequencies) of the aperiodical Xi process is broadly distributed, with notable concentrations in the occipital lobe, temporal lobe, and left parietal lobe (Fig. 4G). The energy of the periodical Rho processes shows a more focal distribution, with high energy localized explicitly in the occipital lobe, corresponding to the Alpha rhythm, and an anterior-to-posterior decrease toward the prefrontal cortex with some energy in the parietal lobes corresponding to the Mu rhythm (Fig. 4H). These energies have been harmonized across recordings to account for differences in EEG amplifiers and site conditions with the Incomplete Observed Linear Mixture Model (IOLMM) model ^[37]^.

Overall, the Gaussianity statistics are relatively low across space for both processes. Non-Gaussianity tests for the Xi process are higher in the frontal region, while the Rho processes have a higher value in the occipital and parietal regions. When observing the maps for the nonlinearity statistic the two processes have distinct spatial distributions. The Xi process is barely considered nonlinear across the entire brain (Fig. 4K). Conversely, the Rho processes show notable nonlinearity in regions that also exhibit high energy, particularly in the parietal lobes (Fig. 4L), and to a lesser degree in the occipital areas.

## 3. Discussion

By analyzing a dataset of 1,771 intracranial EEG (iEEG) and 960 subjects of scalp EEG, we reveal that only a small minority of recordings can be described as both linear and Gaussian. Over 81% of scalp EEG and 67% of intracranial EEG recordings deviate from this ideal, exhibiting significant nonlinearity and/or non-Gaussianity. This challenges the notion that mesoscale brain dynamics are predominantly linear and Gaussian ^[1,2]^. By decomposing the iEEG signals with BiSCA, we find a clear separation between the aperiodic background activity and rhythmic oscillatory components. The broadband Xi process (aperiodic 1/f component) behaves largely as a linear, Gaussian process, whereas the narrowband Rho components (oscillatory peaks, e.g. the alpha rhythm and Mu rhythm) are the primary sources of nonlinearity in the EEG. In other words, the resonant oscillations in the brain carry nearly all of the detected quadratic nonlinear interactions (as evidenced by significant bicoherence), while the aperiodic component shows minimal nonlinear coupling. While previous SPA methods have successfully characterized the power-spectral features of these components, they are blind to phase relationships and therefore cannot assess their nonlinearity.

We observed clear signs of nonlinear resonance phenomena in the data. In the bifrequency domain analysis of the bicoherence, rhythmic peaks (especially the 10 Hz alpha rhythm) exhibited harmonic coupling – for example, frequency-doubling and intermodulation effects were visible as diagonal and off-diagonal bicoherence. These results confirm that well-known cross-frequency coupling phenomena (e.g. harmonics of alpha) are present even in resting-state activity and are quantitatively detectable with our approach. With our BiSCA model we can parameterize the coupling strength between oscillatory peaks.

### 3.1 Linear vs. Nonlinear Modeling Debate

The question of whether large-scale brain signals exhibit nonlinear instead of purely linear behavior is directly answered by our results. As mentioned before, recent studies reporting evidence that macroscopic resting-state dynamics are adequately captured by linear models suggest that microscopic nonlinearities are “averaged out” at the EEG/fMRI level. In contrast, our bichoherence tests revealed that the most EEG channels exhibited higher-order statistical structure. Nonlinear interactions among neural population oscillations seem to persist despite both spatial and temporal averaging. In fact, our finding that a greater percentage of scalp EEG channels appeared nonlinear than iEEG channels suggests that, rather than decreasing nonlinearity as might be expected ^[15,38,39]^, volume conduction primarily spreads this activity across the scalp, contributing nonlinear features to a wide array of channels. Prior work ^[7–10,12,6,13,11]^ showed that nonlinear phenomena like cross-frequency coupling and harmonic generation can occur in neural mass models and actual EEG/LFP recordings. Our findings also suggest that large-scale measures retain higher-order structure reflecting nonlinear resonance in neural circuits. This finding suggests that to capture brain dynamics fully, nonlinear terms are needed. The debate should thus move beyond the binary question of “linear vs. nonlinear models” and consider how specific nonlinear features (like those we observed in Rho oscillations) can be integrated into interpretable models of brain activity. It would therefore be premature to assert the superiority of linear models ^[1]^ without optimally tuning and evaluating nonlinear models on an equal footing.

### 3.2 Interpretation of Oscillatory vs. Aperiodic Components

The biophysical separation ^[40]^ of components in BiSCA follows the assumption that two processes are additive and independent, deriving from different layers of the cortex ^[41–45]^. A recent study provides further neurophysiological evidence for biophysical separation, further demonstrating with biophysical modeling and experimental evidence that the Xi process is a genuine signal generated by arrhythmic network activity, rather than merely an artifact of overlaid brain rhythms ^[33]^.

The analytical separation of components in BiSCA is theoretically grounded in two principles. First, for statistically independent processes, the cumulant spectrum (including the bispectrum) is additive ^[27]^. Second, and more broadly, even for single nonlinear systems described by Volterra functional series ^[46,47]^, the bispectrum can be expressed as a sum of additive frequency components. Each component is analytically defined by the system’s linear and quadratic transfer functions, allowing for the formal description and identification of nonlinear interactions.

A central finding of our study is that the aperiodic (Xi, ξ) component of the EEG signal—which is widely considered to exhibit a 1/f-like power law—is predominantly a linear and Gaussian stochastic process. At first glance, this observation may appear to challenge established opinions that the aperiodic component has a 1/f spectrum. If true, this would tie the Xi process to critical, avalanche-like processes. We have not been able to find convincing reports in the literature that the aperiodic component is a 1/f process. On the contrary, in our findings, the Xi process showed negligible bicoherence and near-Gaussian amplitude statistics across the cortex. This suggests that the aperiodic background can be modeled as a superposition of many independent or weakly interacting sources (consistent with a linear Gaussian process). The linearity of the Xi component can thus be understood as an emergent property, likely arising from the superposition of numerous weakly correlated neural sources where nonlinearities are averaged out at the macroscopic scale. This empirical finding is supported by recent theoretical models such as that of Kramer & Chu, (2024)^[34]^, who demonstrated that a general, noise-driven dynamical system, when linearized around a stable equilibrium, naturally generates spectra with 1/f-like characteristics. Early research by this group (Valdes 1990 et al.)^[48,49]^ on the spatial structure of the Xi process used frequency domain tests and found that the aperiodic cross-spectrum matched a brain eigenmode pattern characteristic of an isotropic random process. It was proposed that such a process may result from short-range cortical neural interactions with exponential decay. This provides a framework for understanding the Xi process throughout the cortex. This mechanism is conceptually identical to a basic linear filter in control theory, whose characteristic Bode plot slopes produce such power-law spectra, offering a parsimonious explanation. It provides a compelling theoretical foundation for why a macroscopically linear stochastic process can produce the observed aperiodic background without invoking complex nonlinear mechanisms. It is also crucial to consider the specificity of our methodology; the bicoherence is a powerful tool for detecting quadratic nonlinearity, and our results show a definitive lack of such second-order interactions in the Xi component. While we cannot formally exclude other, more complex forms of nonlinearity, our work robustly refines the debate on neural linearity.

In contrast, the Rho components (distinct spectral peaks corresponding to oscillatory neural generators) display robust nonlinear signatures. For the alpha rhythm (around 10 Hz) and other peaks, we found significant bicoherence indicating phase-locked interactions, such as harmonics (e.g. 20 Hz, 30 Hz components related to alpha) and possible subharmonics or cross-frequency coupling with other bands. The fact that the full EEG signal’s nonlinearity closely tracked that of the Rho component (and not Xi) implies that oscillatory networks are the location of nonlinear resonant dynamics. The resonances detected seem to correspond mainly to thalamocortical loops in posterior cortices for alpha, and sensorimotor cortices for mu. This is also consistent with corticothalamic neural field models, which generate bursts of nonlinear alpha oscillations through noise-driven exploration of a subcritical Hopf bifurcation ^[50]^. The Valdes et al. (1990)^[48,49]^ paper found that spatial isotropy was not present at the frequencies corresponding to the spectral peaks, supporting the conclusion that nonlinear resonances are more locally distributed on the cortex.

#### Dissociation Between Power and Nonlinearity

An interesting observation is the mismatch between the cortical distributions of oscillatory power and degree of nonlinearity. This spatial dissociation suggests that a high-power spectrum of oscillation does not relate systematically to strong nonlinearity. Consider the posterior alpha rhythm, with high power but weaker nonlinearity than the mu rhythm.

There is evidence that in usual recordings of thrytum, there is a mixture of intermittent, high-amplitude bursts of nonlinear activity interspersed with periods of low-amplitude, quasi-linear dynamics close to a stable fixed point ^[51]^. Such bistable dynamics can emerge from models near a subcritical Hopf bifurcation ^[50,52]^, would yield high spectral power due to the amplitude of the bursts, but the bicoherence, when averaged over a long time window, would be diluted by the intervening linear periods. In contrast, the less powerful parietal mu rhythm may operate in a more persistently nonlinear state, resulting in a disproportionately stronger bicoherence signature. This observation is consistent with a prior study^[53]^, which reported that there are power peaks that coexist in the Alpha and Beta bands in the parietal lobe, while only peaks in the Alpha band appear in the occipital lobe. Our BiSCA method allows us to specifically pinpoint functionally distinct sources of nonlinearity. a similar EEG source-reconstruction study ^[54]^ also describes this dissociation between the spatial patterns of power and bicoherence. It seems that different neural circuits and networks have different dynamical regimes which are projected onto the statistical signatures at the EEG. Some thalamic circuits operate closer to linearity and have high power, while others operate more nonlinearly but have lower power.

### 3.3 Methodological Limitations

#### Spatial Sampling and Data Considerations

The scalp EEG dataset we used includes recordings from 960 healthy multinational subjects and iEEG samples comprised by 1,771 channels from 106 patients. the nature of this data should be considered. The iEEG recordings come from postoperative epilepsy patients, which might have influenced the prevalence or distribution of nonlinear dynamics (e.g., due to cortical irritability or medication effects). However, we observed consistent patterns between iEEG and healthy EEG. In addition, these patterns are also consistent with a prior paper with healthy subjects^[54]^, suggesting our conclusions are not driven by covert pathological activity. Another consideration is the volume conduction and presence of a reference in scalp EEG, which can smear true source interactions. We mitigated this by focusing many analyses on iEEG (where signals are more focal) and by using a conservative statistical threshold (FDR q=0.001). Still, some spurious mixing in scalp recordings cannot be entirely ruled out. To resolve these issues, source localization combined with BiSCA will help to confirm that the identified nonlinear interactions correspond to true neurophysiological coupling rather than mixing artifacts.

### 3.4 Implications

#### Nonlinear Resonance as an Organizing Principle

We found that the dominant EEG exhibits nonlinearity. This finding agrees with copious theoretical work that models these oscillations with nonlinear models. Explicitly linking the spectral “peaks” to bispectral evidence of coupling, we highlight the importance of nonlinear resonance in its more general formulation. We envisage that our methods may contribute to deal with disparate phenomena like EEG cross-frequency coupling, nested oscillations (e.g., phase–amplitude coupling), and event-related harmonic responses within a unified resonance-based theory. It also supports methods like ICA or nonlinear signal decompositions, which concentrate on the nonlinear and non-Gaussian aspect of the EEG.

#### Refining Macroscale Models

In practical terms, our findings encourage the development of new macroscale brain models that incorporate nonlinearity with the prior knowledge provided in this paper in a controlled way ^[55–62]^. Such models might treat the aperiodic background as an additive Gaussian noise process (as linear models do) while nonlinear coupling would be reserved for oscillatory subsystems. This hybrid perspective – essentially what our data-driven BiSCA components represent – could improve how we simulate or predict EEG dynamics. For example, large-scale neural mass or neural field models might include nonlinear feedback only in certain frequency-generating circuits (like alpha or mu generators), which could reproduce the selective nonlinearity we observed. BiSCA or methods like it might provide data-driven statistics that neural models should match.

#### Extend the BiSCA to nonstationary EEG/iEEG

BiSCA analyzes the system’s quadratic nonlinearity (bispectrum/bicoherence) based on the assumption of stationarity. This is a condition that the providers of the data tried to ensure. Nevertheless, there is no question that brain activity is nonstationary. It is clear that phenomena such as multistability indicate dynamic changes in the nonlinear properties ^[50,51,63]^ in EEG data, which shows that a linear system with a single global attractor is an insufficient model. The BiSCA approach is a tool that assumes a stationarity complementary perspective. We must extend BISCA to identify persistent quadratic interactions from those that change dynamically ^[64]^.

## 4. Materials and Methods

### 4.1 Materials

We analyzed resting state EEG data from 960 healthy participants drawn from the globally diverse HarMNqEEG dataset ^[65]^. Participants were selected from normative or control groups, excluding those with neurological/psychiatric conditions. EEG recordings consisted of eyes-closed resting-state data. For all subjects, 18 channels were included (excluding Pz) from the 10-20 system after average referencing. Samples recorded for less than 1 minute, such as the Medicid-4-Cuba-2003, were excluded. Preprocessing involved converting signals to an average reference, segmenting data into 300 time points (1.5 seconds), and overlapping segments to have 157 segments per channel to maintain consistency with the iEEG data length and segmentation. Segments were retained only if artifact-free, with no interictal discharges or slow-wave anomalies.

For Resting-state Intracranial EEG (iEEG), we used the dataset of eyes-closed resting-state wakefulness recordings, sEEG/ECoG recordings from the Montreal Neurological Institute iEEG atlas ^[53]^. These iEEGs were recorded from 106 patients with refractory focal epilepsy (1,772 electrodes), ensuring that tracings retained were from non-lesional, non-seizure onset zones, free of interictal discharges, slow-wave anomalies, or post-seizure/stimulation effects. Preprocessing involved bandpass filtering (0.5–80 Hz), down-sampling to 200 Hz, and removing zero values. The study focused on 60-second eyes, in order to be consistent with the EEG data set.

### 4.2 Method

#### 4.2.1 Multitaper Estimator for the Spectrum

Each segment of the signal was subjected to a Discrete Fourier transform 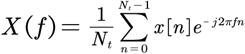, where *N*_*t*_, is the number of time points for each epoch. To balance the frequency resolution and computational resources, in our study, the window size was 300 time points (1.5 seconds) segments. To estimate the power spectrum, S(*f*) = E [*X*(*f*)*X* ^*^(*f*)] we used overlapped segments by 75% and also applied the Multitaper method, which creates windowed versions of the original time signal *x*_*k*_ (*n*) = *x*(*n*)*v*_*k*_(*n*) ^[66,67]^ in order to reduce frequency leakage bias and increase the consistency of the spectral estimate, *k* is the index of taper. For this purpose, we used the Sine tapers *v*_*k*_ ^[68]^ to diminish amplitude bias further. The resulting estimation procedure is then:

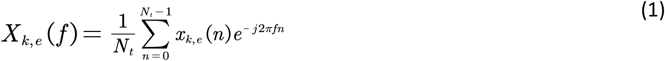

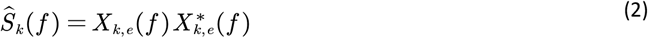

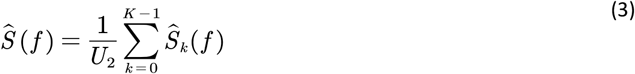

where *U*_2_ =*K*/*N*_*t*_. In this paper, we use *NW* = 1.5, i.e. *K* = 2*NW* − 1 = 4 for the recordings. *NW* This critical parameter controls the estimator’s trade-off between frequency resolution and variance.

#### 4.2.2 Multitaper Estimator for the Bispectrum

We apply the theory developed by Brillinger for nth order cumulant spectra ^[69,70]^. The usual power spectrum is the 2nd-order cumulant spectrum we just presented above. In this paper, we also deal with the bispectrum, which is the 3rd-order cumulant spectrum, a function of the two frequencies (bifrequency) *f*_1_ and *f*_2_

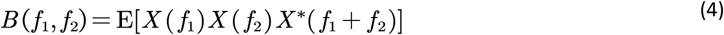

As mentioned above, the bispectrum has the favorable properties of zeroing Gaussian processes, being constant for linear systems with non-Gaussian input, and reflecting quadratic nonlinear interactions between bifrequency pairs *f*_1_ and *f*_2_ and their sum *f*_1_ + *f*_2_.

The Multitaper estimator for the bispectrum and bicoherence ^[71,72]^

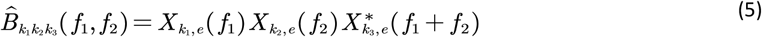

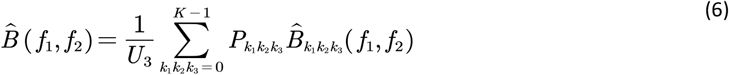

To preserve the correct scale, the ‘bienergy’ 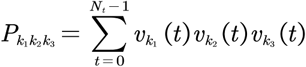 and 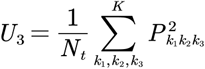.

We estimate the bicoherence *b*(*f*_1_*f*_2_) using the following normalization of the bispectrum ^[73]^, which benefits the statistical tests defined in the next section:

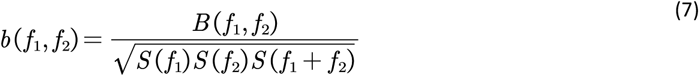

#### 4.2.3 Test for Gaussianity and linearity through Higher-Order Spectrum

To characterize the underlying dynamics of the time-series data, we employed higher-order spectral analysis to test for linearity and Gaussianity. Linear and nonlinear systems are fundamentally distinguished by their properties of response to input frequencies: a linear system processes frequencies independently, whereas a nonlinear system generates new harmonic and intermodulation components through frequency interactions. Bicoherence is utilized to detect and quantify this quadratic phase coupling.

The theoretical foundation for this method is predicated on the behavior of the bicoherence for a linear time-invariant (LTI) system. For an LTI system driven by a non-Gaussian input, the magnitude of the bicoherence, |*b*(*f*1,*f*_2_)|, is constant across all frequency pairs and is determined solely by the input signal’s third-order 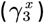 and second-order 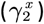 cumulants ^[74]^:

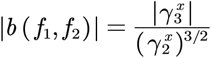

This property yields a clear framework for hypothesis testing, used in Table 1:

1. A **Gaussian Linear (GL) process** is the null case. It has a third-order cumulant of zero, resulting in a bicoherence that is identically zero across all frequencies.
2. A **Non-Gaussian Linear (NGL) process** is driven by a non-Gaussian input passing through a linear system. This results in a **diffuse, widespread elevation** of bicoherence across the entire frequency plane.
3. A **Gaussian Nonlinear (GNL) process** involves a Gaussian input passing through a nonlinear system. The nonlinearity generates phase coupling at specific frequency interactions, producing **sharp, localized peaks** in the bicoherence plot against a near-zero background.
4. A **Non-Gaussian Nonlinear (NGNL) process** represents the combined case where a non-Gaussian input drives a nonlinear system. Its bicoherence exhibits both localized peaks superimposed on a diffuse, non-zero background.

To achieve a more nuanced understanding of underlying dynamics and explicitly separate system nonlinearity from input non-Gaussianity, this refined framework tries to detail the nonlinear case to GNL and NGNL. The test for intrinsic non-Gaussian input assesses the diffuse, central tendency of bicoherence, whereas the test for system nonlinearity targets sparse, high-magnitude peaks indicative of quadratic phase coupling. It is important to note that the term “non-Gaussian” within this framework refers to any deviation from Gaussianity in the observed time series that is not solely a result of system-induced phase coupling. This includes both non-Gaussian properties of the system’s innovations (inputs) and the presence of independent additive non-Gaussian noise, both of which manifest as a diffuse, non-zero bicoherence.

The statistical implementation of these tests relies on the asymptotic distribution of the bispectrum estimator 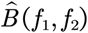. For a zero-mean stationary process, the bispectrum asymptotically follows the complex Gaussian distribution, its expected value and variance are given ^[69]^

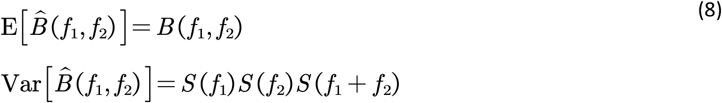

Statistical tests for (quadratic) Gaussianity and linearity can be effectively formulated using bicoherence, which measures quadratic phase coupling. The traditional test proposed by ^[36]^ relies on the sum of squared bicoherence values, comparing the result to a chi-squared distribution. However, this sum-based approach is overly sensitive to extreme values, which can arise from strong, localized nonlinear interactions like narrow-band phase coupling. This sensitivity can cause the test to reject the null hypothesis of Gaussianity even when the process only exhibits isolated nonlinearity, leading to false positives. The sum is easily dominated by a few large values, misrepresenting the overall distribution as non-Gaussian.

For testing linearity specifically, traditional methods like the interquartile range (IQR) test are often inadequate. The IQR focuses on the central 50% of the bicoherence distribution, making it a poor detector of narrow-band quadratic phase coupling. Such interactions manifest as sparse, extreme values in the bicoherence map, which have little impact on the IQR. Consequently, the test lacks the power to identify significant but isolated nonlinear peaks.

To address these limitations, we propose two robust tests that enhance the Hinich framework by targeting distinct aspects of deviation from the null hypothesis. The first is a median-based test to assess central tendency for Gaussianity, and the second is a maximum-based test to detect nonlinear interactions. Both tests leverage bicoherence’s ability to measure phase coupling, with their statistical properties validated through the theoretical derivations below.

##### 4.2.3.1 Median-Based Test for Gaussianity

To provide a robust assessment of Gaussianity that is less affected by outliers, we introduce a test focused on the median of the squared bicoherence values. We first define a normalized statistic, *b*^(*G*)^, which represents the median bicoherence magnitude and serves as a measure of effect size for asymmetricity:

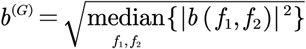

The formal test statistic, *t*^(*G*)^, is a scaled version of this quantity:

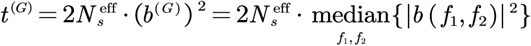

where 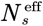 is the effective number of segments, and the median is computed over all bifrequency pairs (*f*_1_*f*_2_). Under the null hypothesis of Gaussianity and linearity, this statistic follows an asymptotic normal distribution. The test evaluates:

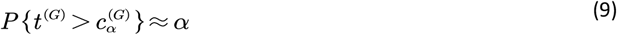

Where 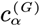 is the critical value from the asymptotic normal distribution at the significance level *α*. This test excels at detecting non-Gaussianity.

This test is derived from the asymptotical normal distribution of bicoherence. Under the null hypothesis of Gaussianity, for each independent bifrequency pair, the scaled squared bicoherence 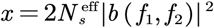 follows a chi-squared distribution with two degrees of freedom *x* ~ *χ*^2^(2). The test statistic *t*^(*G*)^ is the sample median of *P* such independent variables. For a large number of bifrequency pairs *P*, the sample median is asymptotically normal. The population median of a *χ*^2^(2) distribution is *m* =2ln2 ≈ 1.386. This is the expected value of our test statistic, *E* [*t*^*(G)*^] ≈2ln2. The asymptotic variance of a sample median from a sample of size *P* is given by 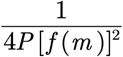, where *f*(*m*) is the value of the probability density function (PDF) at the population median *m*. The PDF of a *χ*^2^(2) distribution is 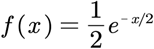. At the median *m* = 2ln2, the PDF value is 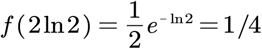. Therefore, the variance is:

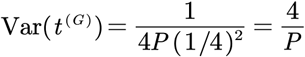

Thus, the test statistics follow the asymptotic distribution:

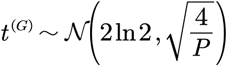

##### 4.2.3.2 Maximum-Based Test for Nonlinearity

To identify nonlinear interactions that may be missed by the median test, we introduce a test based on the maximum squared bicoherence value. The normalized statistic, *b*^(*L*)^, directly captures the magnitude of the strongest quadratic phase coupling:

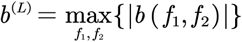

The corresponding test statistic, *t*^(*L*)^, is its scaled squared value:

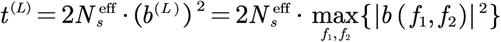

To distinguish localized linearity from background non-Gaussianity, we estimate a non-centrality parameter 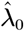. The distribution of *t*^(*L*)^ is modeled using extreme value theory, specifically a Gumbel distribution. The test evaluates:

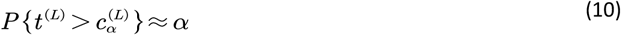

Where 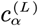 is the critical value from the Gumbel distribution at the significance level *α*. This test is designed to pinpoint significant localized nonlinearity.

To account for baseline non-Gaussianity, we estimate a non-centrality parameter *λ*_0_:

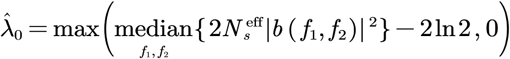

This reflects that the median of a non-central *χ*^2^(2, *λ*_0_) distribution is approximately 2ln + *λ*_0_ for small *λ*_0_. Using extreme value theory (EVT), the maximum of *P* independent *χ*^2^(2, *λ*_0_) variables is approximated by a Gumbel distribution. The normalizing constants are:, *b*_*p*_ = *F1*^−1^(1−1/*P)*, the value where the cumulative distribution function (CDF) *F* of *χ*^2^(2, *λ*_0_) Reaches 1 − 1 / *P*. And 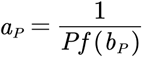, where *f* is the PDF of*χ*^2^(2, *λ*_0_). Thus, the nonlinearity test statistic is distributed as:

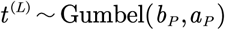

##### 4.2.3.3 Estimation of the Effective Number of Segments

The theoretical distributions described above assume that bicoherence values are derived from independent data segments. However, modern spectral estimation techniques like the multitaper method we used here and the use of overlapping segments introduce dependencies. This means the nominal number of segments, *N*_*s*_, is larger than the effective number of independent segments, 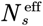. Using the nominal *N*_*s*_ in the test statistics will lead to incorrect scaling, thresholds, and p-values.

To ensure accuracy, we estimate 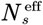 with the following procedure:

1. Fit an AR model: Fit an autoregressive (AR) model to the observed time series *x*(*t*) using the Burg method. Select the optimal model order using the Corrected Akaike Information Criterion (AICc).
2. Generate surrogate data: Generate *M* surrogate time series (e.g., *M* = 100) using the fitted AR model with Gaussian white noise as innovations. Each series should have the same length *N* as the original data.
3. Compute bicoherence: For each of the *M* surrogate series, compute the bicoherence *b*(*f*_1_,*f*_2_) using the exact same parameters (e.g., Multitaper settings, window length, overlap) as used for the original data.
4. Calculate the mean squared bicoherence: For each surrogate series, calculate the mean of the squared bicoherence magnitudes over all *P* valid bifrequency pairs:

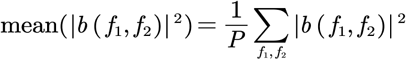
5. Estimate 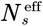: Average the result from step 4 across all *M* simulations. Under the null hypothesis, 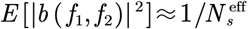 Therefore, the effective number of segments can be estimated as:

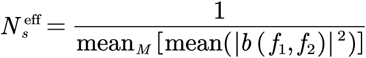

This estimated 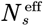 should replace *N*_*s*_ in all calculations for the *t*^(*G*)^ and test statistics and their corresponding distributions, thereby accounting for dependencies and ensuring the validity of the statistical tests.

#### 4.2.4 BiSpectral EEG Component Analysis (BiSCA)

##### 4.2.4.1 Whole model

After obtaining the empirical and higher-order spectra, this paper proposed a BiSCA model specifically designed to detect nonlinear resonance by jointly parametrizing the spectrum and bispectrum. This combined model approach is essential for identifying nonlinear resonance phenomena that cannot be detected by traditional spectral analysis alone. Here, the Xi trend and Rho peaks (including Alpha and all other oscillatory peaks) are modeled with two types of independent processes, allowing them to be additive on the natural scale. Additionally, the analysis includes both the spectrum and the bispectrum to understand the components comprehensively.

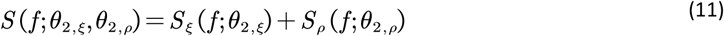

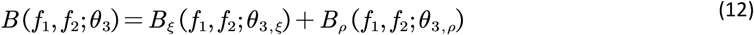

where different *θ* are the parameters for different components in each order.

##### 4.2.4.2 Model for Spectrum

In this paper, we put all peaks in the group of Rho processes for several reasons. 1) To be consistent with the conventional XiAlpha model, we model the trend as the Xi process and the oscillatory rhythms as the Alpha process. 2) The models of oscillation peaks have similar structures. 3) We focus on univariate analysis, which views the neural masses as a single integrated oscillator. For the spectra, each component consists of a sum of squares of kernel functions. Since here we use the transfer function to connect different orders of spectra, the sum of the squares of the frequency transfer function of each component approximates the modeled theoretical power spectra here

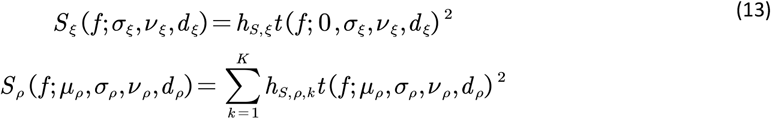

The kernel function models the oscillatory rhythm peaks in the spectrum domain. The *h* is the scale parameter for each peak, and the peak number *K* represents the number of harmonics to be modeled. We offered two strategies for determining the center frequency of peaks: (1) the {*μ* _*ρ,k*_} is 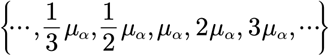, in practice, the order *k* is chosen with the Akaike information criteria. This circumstance models the alpha rhythm with the simplest condition that the resting state EEG is the output of the system excited with Gaussian white innovation, and the 1st-order transfer function has a single tone at Alpha; the rest of the peaks in other bands are the harmonics of the fundamental oscillation peak. (2) fit each of the {*μ* _*α,k*_} through the spectra and then adjust with the joint fitting with the bispectrum. Option (1) offers frequency peak organization following the conventional broadband analysis; option (2) will fit the components better but is more complicated. This paper demonstrates the result with option (2), which is more general and can use the parameter for bicoherence to interpolate the phase relationship between the peaks. The goodness of both option fit is shown in the supplementary.

The student t kernel function has been used in previous studies of spectrum fitting ^[75–77]^. Here, we modified the student t function to model the amplitude of the 1st-order frequency transfer function, which also extends the adjustable heavy-tail version of the Lorentzian (Cauchy-Lorentz distribution function).

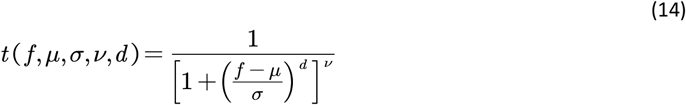

The parameter *σ* controls the width of the kernel function. The parameter *d* controls the flat top of the kernel since the Multitaper method is smooth in the frequency domain and distorts the shape function; in practice, the value is larger than 2. The parameter *ν* controls the kurtosis or the heavy tail of the peak. Special case: *d* =2 it turns to the student t function; *v* = 1,*d* = 2 it turns to the Lorentzian kernel.

##### 4.2.4.3 Model for bispectrum

To simplify the modeling process, we did not directly use the theoretical composition form of transfer function kernels to each order of the spectrum ^[46,47,78–80]^. Instead, we focus on capturing the primary relationships of position and order by parameterizing the theoretical bispectrum *M*_3_(*f*_1_,*f*_2_) as follows:

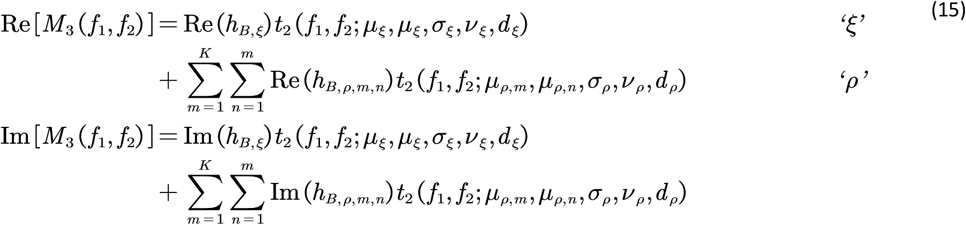

This formula captures the essential features of the bispectral peaks without resorting to complex transfer function compositions. The BiSCA models the real and imaginary parts of the bispectrum separately since the phase information of the bispectrum is necessary for the discrimination of systems ^[81]^. The kernel function for the bispectrum is

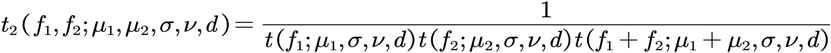

##### 4.2.4.4 Likelihood

The likelihood of the model follows the asymptotically Gaussian distribution of the log-scale spectrum ^[82]^ and natural scale bispectrum^[83]^.

To ensure that the Xi process exhibits a wide spectrum, we add a likelihood term for the Xi process to encourage fits that adhere to the overall trend. The joint likelihood for the full model is:

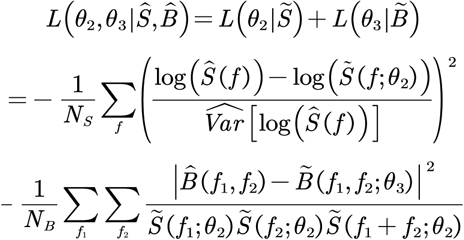

where the *Ŝ*(*f*) and 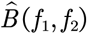 is the Multitaper power spectrum and bispectrum estimate by formula (3) and (6), 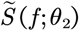 and 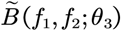 is the modeled theoretical power spectrum and bispectrum.

This likelihood-based method makes it advantageous to use statistical methods to do the model comparison.

##### 4.2.4.5 Fit with the Levenberg-Marquardt method

We estimate the model parameters using the Levenberg-Marquardt (LM) optimization algorithm. The LM method is a standard technique for solving nonlinear least squares problems, effectively balancing the Gauss-Newton algorithm and gradient descent. It is particularly suitable when the objective function is the sum of squares of loss functions, as in our joint likelihood estimation, which is also a heteroscedastic curve/surface fitting problem. Employing the LM method, we iteratively update the parameter estimates to minimize the discrepancy between the empirical and theoretical spectrum and bispectrum, ensuring convergence to an optimal solution. The LM algorithm is not a global optimization method. Therefore, we fit the spectra parameters first and then share the parameters with the bispectrum as a warmup for the joint fit.

##### 4.2.4.6 Bicoherence for each component

We can then obtain the bicoherence for each component

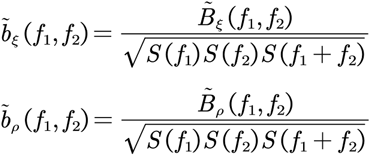

We use full variance to normalize the component bicoherence to avoid instability and ensure the additive way to model the bicoherence.

##### 4.2.4.7 Test for components

Then we recall the test (9) and (10) for testing the non-Gaussianity and linearity of each component.

## 6. Data availability

The scalp EEG and intracranial EEG (iEEG) data analyzed in this study were obtained from the work of Li et al., (2022)^[65]^ and Frauscher et al. (2018)^[53]^, respectively. Ethical approval was not required for this specific study as it exclusively analyzed pre-existing, publicly available, and anonymized datasets (the HarMNqEEG dataset and the Montreal Neurological Institute iEEG atlas). The collection of the original data received ethical approval from the respective local authorities, as detailed in the original publications ^[53,65]^.

## 7. Code availability

The analysis code used in this study is publicly available on GitHub at https://github.com/rigelfalcon/BiSCA.

## 8. Acknowledgement

We thank Dr. Nicolás von Ellenrieder for his invaluable assistance in providing the MNI iEEG Atlas dataset. This work was supported in part by the STI 2030-Major Projects (Grant No. 2022ZD0208500), the National Natural Science Foundation of China (Grant No. W2411084), the Key Research and Development Projects of the Science and Technology Department of Chengdu (Grant No. 2024-YF08-00072-GX), the CNS Program of the University of Electronic Science and Technology of China (UESTC) (Grant No. Y0301902610100201), and the Chengdu Science and Technology Bureau Program (Grant No. 2022-GH02-00042-HZ). L.M. gratefully acknowledges personal support from the Hundred Talents Program of UESTC, the Outstanding Young Talents Program (Overseas) of the National Natural Science Foundation of China, and talent programs of Sichuan Province and Chengdu Municipality. M.B. acknowledges funding from the National Health and Medical Research Council (NHMRC, Grant No.

APP2008612).

## Supplementary Materials

### S.1 Mathematical notations

**Table S1.**
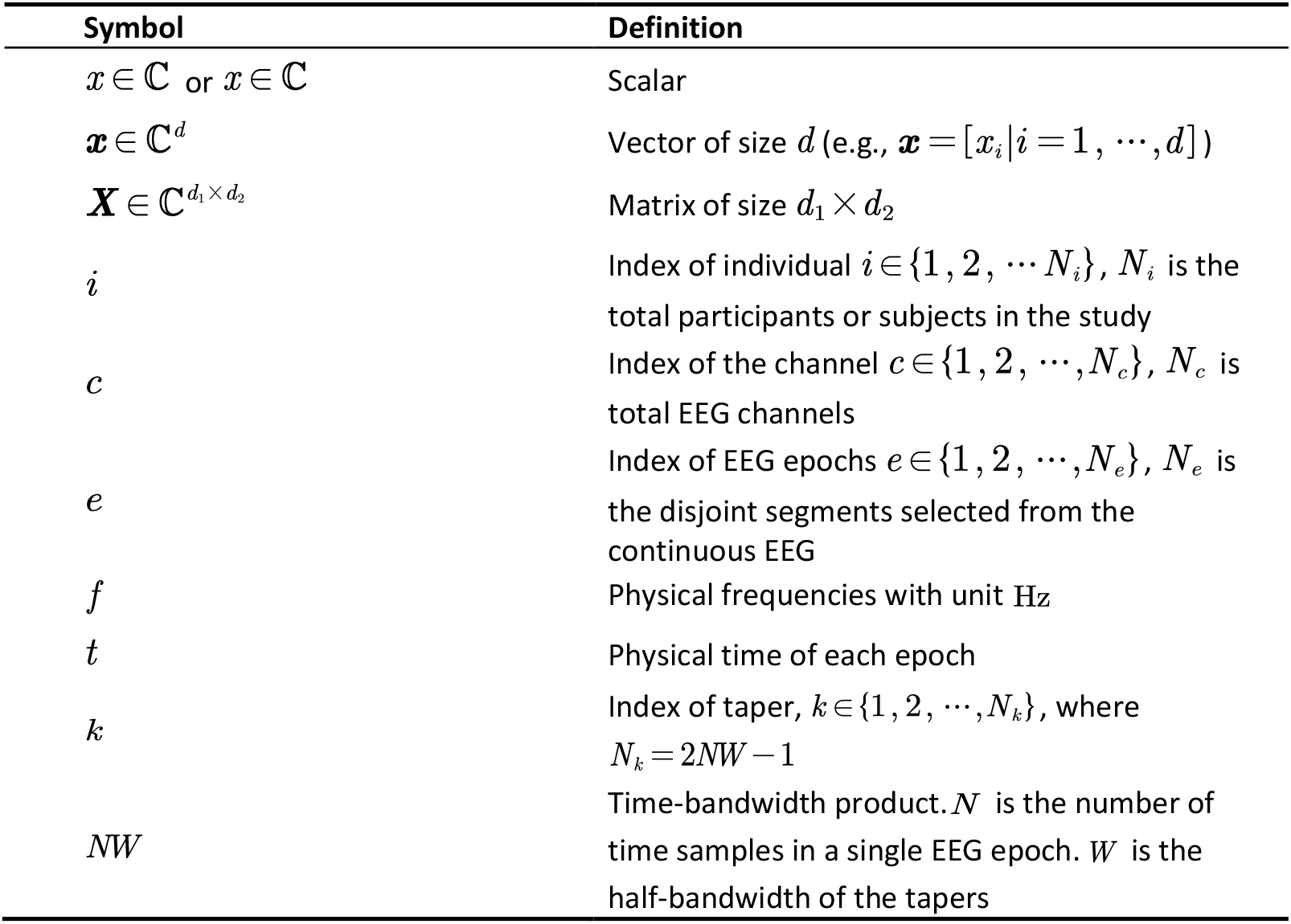
Summary of symbols.

| Symbol | Definition |
| --- | --- |
| $x \in \mathbb{C}$ or $x \in \mathbb{C}$ | Scalar |
| $\mathbf{x} \in \mathbb{C}^d$ | Vector of size $d$ (e.g., $\mathbf{x} = [x_i \vert i = 1, \dots, d]$) |
| $\mathbf{X} \in \mathbb{C}^{d_1 \times d_2}$ | Matrix of size $d_1 \times d_2$ |
| $i$ | Index of individual $i \in \{1, 2, \dots, N_i\}$, $N_i$ is the total participants or subjects in the study |
| $c$ | Index of the channel $c \in \{1, 2, \dots, N_c\}$, $N_c$ is total EEG channels |
| $e$ | Index of EEG epochs $e \in \{1, 2, \dots, N_e\}$, $N_e$ is the disjoint segments selected from the continuous EEG |
| $f$ | Physical frequencies with unit Hz |
| $t$ | Physical time of each epoch |
| $k$ | Index of taper, $k \in \{1, 2, \dots, N_k\}$, where $N_k = 2NW - 1$ |
| $NW$ | Time-bandwidth product. $N$ is the number of time samples in a single EEG epoch. $W$ is the half-bandwidth of the tapers |

**Table S2.** Symbol of quantities and operators.

| Symbol | Name | Definition |
| --- | --- | --- |
| $v_{i,c,e}(t) \in \mathbb{R}$ | EEG potential and its Discrete Fourier transform | EEG potential of the individual $i$, channel $c$ of $e$-th epoch at a time $t$ or frequency $f$ for the Discrete Fourier transform. |
| $v_{i,c,e}(f) \in \mathbb{C}$ | | |
| $S_{i,c}(f) \in \mathbb{R}$ | Theoretical spectra | The theoretical spectrum of an individual $i$ channel $c$ at the frequency $f$ |
| $\hat{S}_{i,c}(f) \in \mathbb{R}$ | Power spectra | The empirical variance vector of $v_{i,c,e}(f)$ across all epochs for an individual $i$ at the frequency $f$ |
| $B_{i,c}(f_1, f_2) \in \mathbb{C}$ | Theoretical Bispectrum | The theoretical bispectrum of an individual channel $c$ at the bifrequency $(f_1, f_2)$ |
| $\hat{B}_{i,c}(f_1, f_2) \in \mathbb{C}$ | Empirical Bispectrum | The empirical bispectrum of process $v_{i,c,e}(t)$ across all epochs for an individual $i$ channel $c$ at the bifrequency $(f_1, f_2)$ |
| $b_{i,c}(f_1, f_2) \in \mathbb{C}$ | Population Bicoherence | Population normalized bispectrum for individual $i$ and channel $c$ at the bifrequency $(f_1, f_2)$ |
| $\hat{b}_{i,c}(f_1, f_2) \in \mathbb{C}$ | Multitaper Bicoherence | Multitaper bicoherence estimator for individual $i$ and channel $c$ for bifrequency $(f_1, f_2)$ |
| $\tilde{b}_{i,c}(f_1, f_2) \in \mathbb{C}$ | BisCA Bicoherence | BiSCA -model estimator for individual $i$ and channel $c$ for bifrequency $(f_1, f_2)$ |
| $\mathbb{E}[\cdot]$ | Expectation | The expectation of a random variable |
| $\text{Var}[\cdot]$ | Variance | The variance of a random variable |
| $[\cdot]^*$ | Complex conjugate | The complex conjugate of a variable |

### S.2 Summary of EEG Spectral Parameterization Methods

As a typical machine learning problem, many toolboxes implemented the curve fitting for spectroscopic data analysis — such as SpectroChemPy ^[84]^, PNNL Chemometric Toolbox^[85]^ astronomy data modeling ^[86]^, SLM (Shape Language Modeling) ^[87]^, LMFIT (Levenberg-Marquardt least-squares minimization) ^[76]^. This paper emphasizes studies that have been applied specifically to EEG spectral analysis with certain constraints of biophysical prior.

**Table S3.**
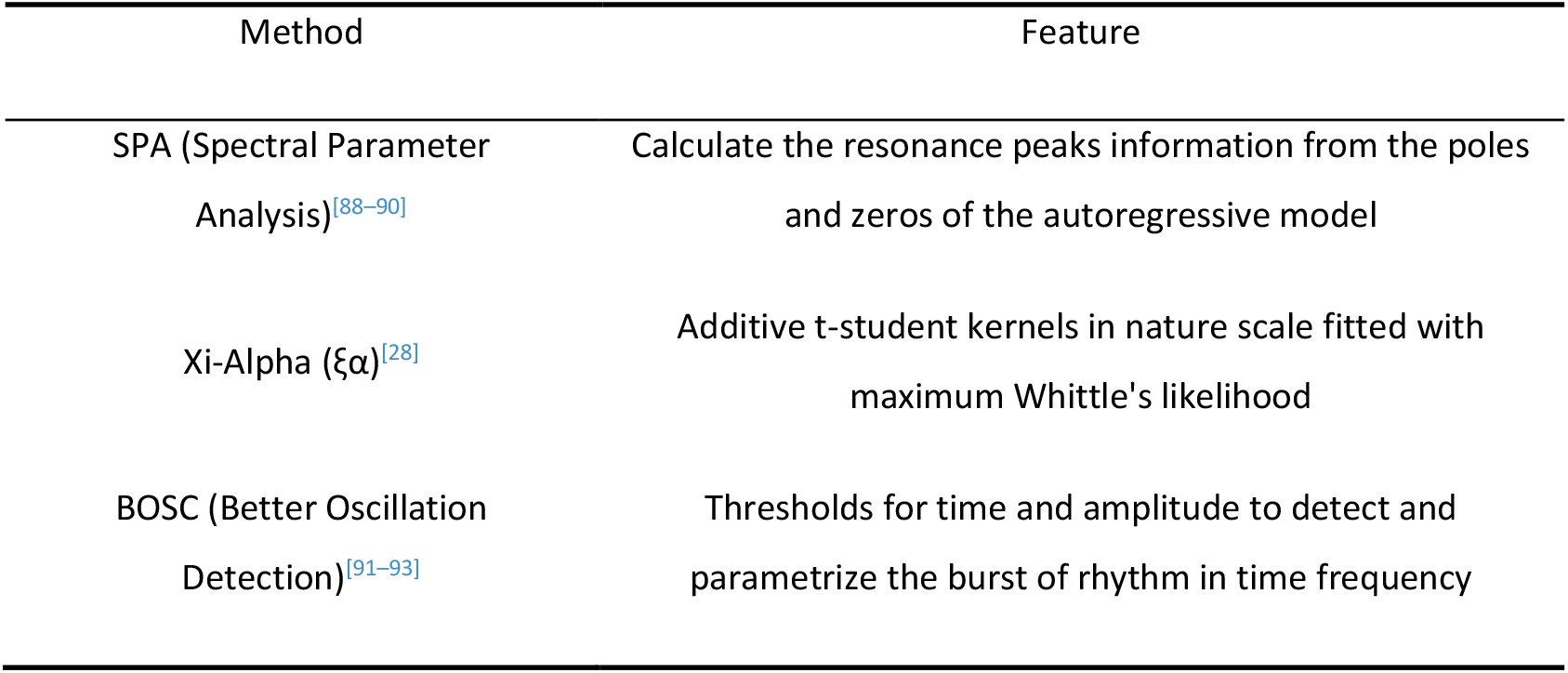
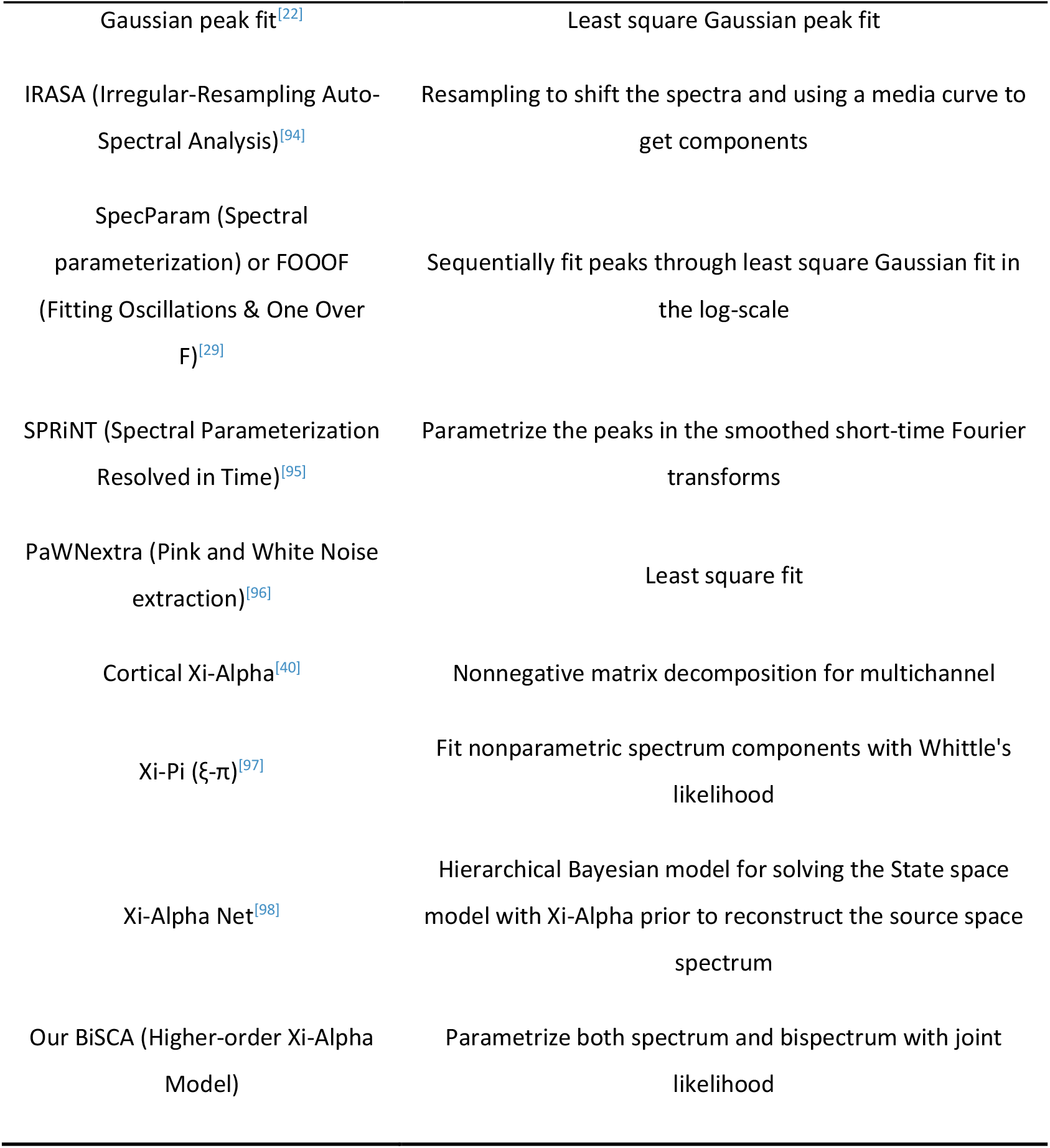
Summary of EEG Spectral Parameterization Methods.

### S.3 Simulation

**Fig. S1.**
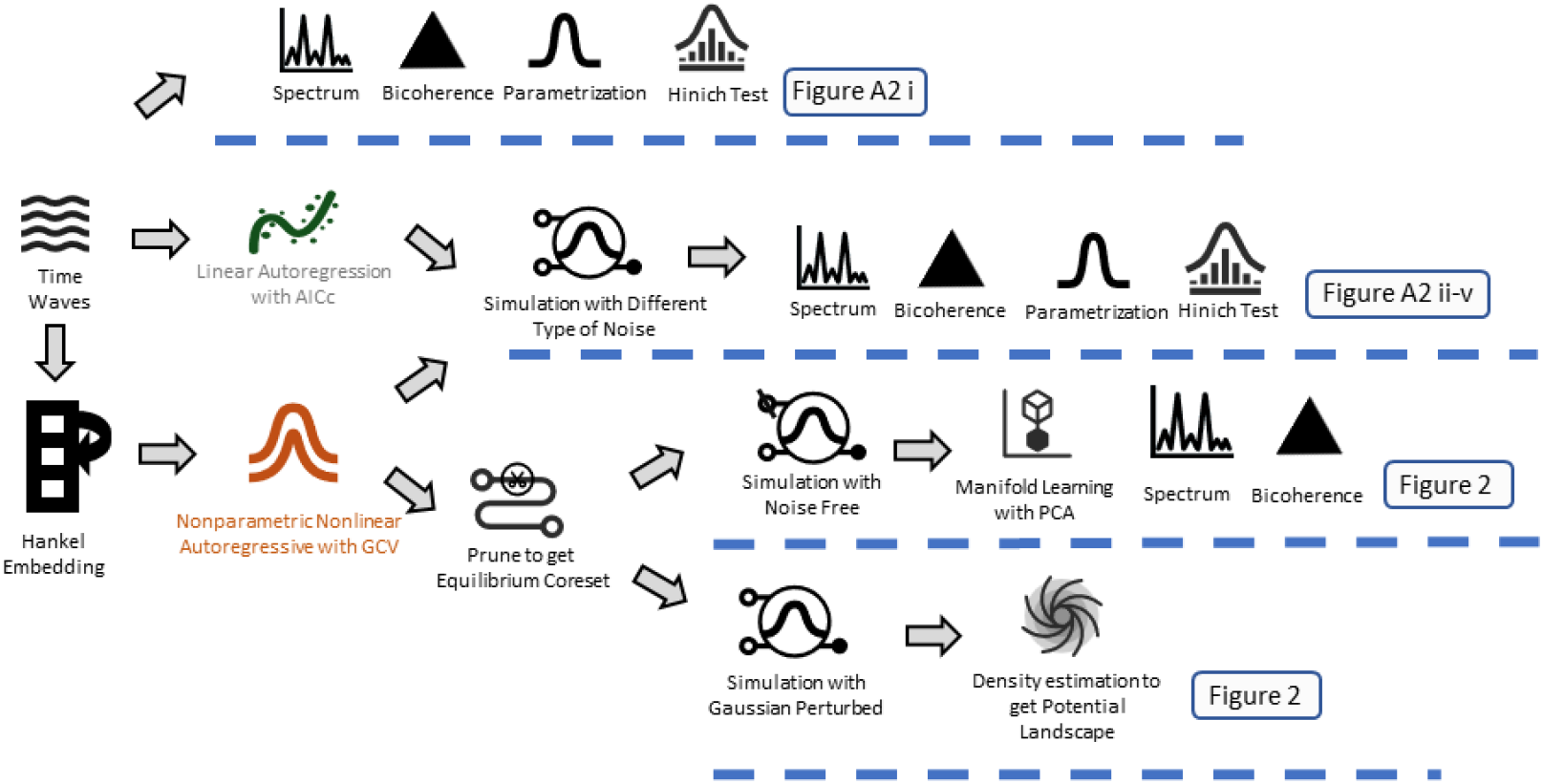
Flowchart of the Simulation and Analysis Pipeline. This diagram illustrates the methodology for analyzing real intracranial EEG (iEEG) data and generating four corresponding types of synthetic surrogate data to test for nonlinearity and non-Gaussianity. (1) Direct Analysis of Real Data (Top Path, Ref: Fig. S2 i): The process begins with a real iEEG time series. This signal is directly subjected to spectral and bispectral analysis to compute its empirical power spectrum and bicoherence as a real-data baseline. (2) Linear and Nonlinear Surrogate Generation (Middle Path, Ref: Fig.S2 ii-v): A linear Autoregressive (AR) and a Nonparametric Nonlinear Autoregressive (NNAR) model is fitted to the real iEEG data. The optimal model order is selected using the Akaike Information Criterion corrected (AICc). This AR model and NNAR model are then used to generate two types of time series by driving it with different stochastic innovations: Linear/Nonlinear Gaussian: Using Gaussian (normal) distributed noise. Linear/Nonlinear Non-Gaussian: Using skewed, non-Gaussian noise (Pearson Type III distribution). (3). Simulation demonstration of the bicoherence and time domain Geometric for the Fig.2: after fitting the NNAR model to the real data, only the states close to the equilibrium is be maintained in the model and used to simulate to get spectrum, bicoherence and 2d embedded potential (density) function.

#### S.3.1 Simulation for the Hinich’s test and BiSCA parametrization demonstration

Here is an example of our bicoherence analysis for different types of signals, we put a simulation demonstration consisting of one channel of real data and 4 corresponding synthetic data (linear Gaussian, linear non-gaussian, nonlinear Gaussian, and nonlinear non-gaussian).

1. Real data: one channel from the MNI atlas data, with Fs=200Hz and 12000 time points samples.
2. Linear Gaussian process: This simulation used an Autoregressive (AR) model with Gaussian innovation.

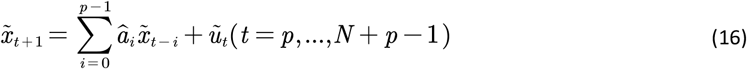

The AR coefficient is fitted with the real data using Burg’s method. In this example, a 45-order AR model is selected with AICc criteria, and the innovation is *N*(0,1). We simulated *N* = 12000 samples with this AR model, also the same length for the rest 3 simulations.
3. Linear non-gaussian process: Instead of using the Gaussian innovation, here we use the Pearson type III distribution^[99]^ as the innovation enables us to control the first to third-order moment for the distribution. The probability density function is

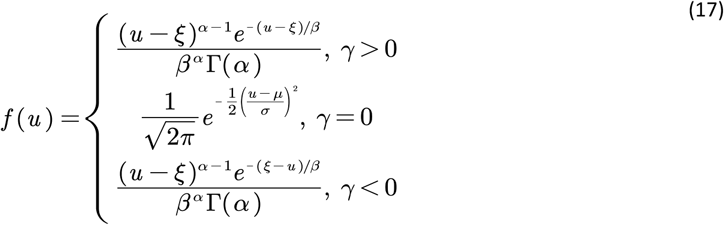

where the parameter of the distribution 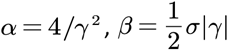, and. The *ξ* = *μ* −2 *σ*/ γ parameter *μ* is the center of distribution, *σ* is the standard deviation, *γ* is the skewness. Here we take *μ* = 0, *σ* = 1 and *γ* = 10.
4. Nonlinear Gaussian process: Instead of linear *P*-lag AR model in the linear case, here we used a nonparametric nonlinear autoregressive (NNAR) model to fit the data ^[100]^, and simulate 12000 timepoints data recursively. Given timeseries 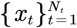, reconstruct phase space via delay embedding:

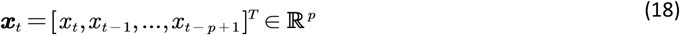

The state evolution is modeled as the discrete nonlinear state space model

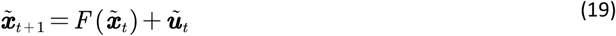

where *F* (·): ℝ^*p*^ → ℝ^*p*^ is a *p* dimensional nonparametric function estimated via adaptive local constant regression (Nadaraya-Waston regression). To simplify the notation to fit regression notation, define embedding vector keys set 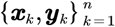 stored all observed data with ***x***_*k*_ = ***x***_*t*_ and, ***y***_*k*_ = ***x***_*t*+1_,+*n*=*N*_*t*_−*p*−1.

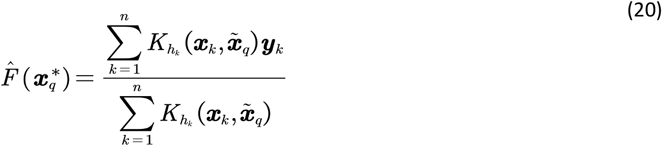

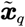 is the query vector state during the N-step forward forecasting. The local weights are calculated by the kernel distance 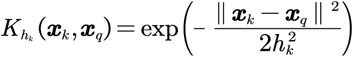 with bandwidth *h*_*k*_. The *h*_*k*_ is the hyper parameter of model, in order to handle non-uniform distribution of the states in the *p* dimension space, we use the *κ*-th nearest neighborhood ***x***_[k, *κ*]_ of each *x*_*k*_ to determine the local bandwidth

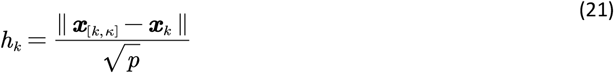

To select *κ*, this paper used a generalized cross-validation (GCV) on the real data 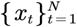 from case 1, to find the best *κ* of and record the corresponding ***h***_*k*_ for each states ***x***_k_ in the one-step forward prediction task.

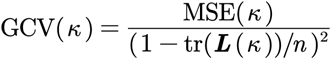

where the smoother matrix with elements

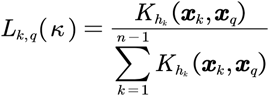

To be computationally efficient, here we use scalar ***h***_*k*_, bandwidth is equal for all dimensions. The embedding dimensions *p* are used the same as linear simulation case 2 and case 3. The nonparametric function 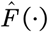 represent all the information of the original (Hankel embedded) state space or the differential manifold of the system, therefore, we didn’t update this nonparametric function during evolution. In the simulation the innovation 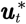 is a multivariable gaussian distribution ***z*** _*t*_ and scaled with covariance from the residual of the one-step forward prediction 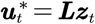, where ***L*** is the Cholesky of the covariance matrix of the residual

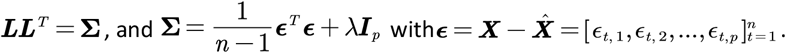

Nonlinear non-gaussian process: The nonlinear system follows the configuration of case 4. Instead of Gaussian innovation, we use Pearson type III (17) to generate the innovation ***z***_*t*_ and then the 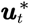.

All 5 cases apply the Multitaper estimation and the BiSCA model fitting with the same parameters as the analysis for real data.

**Fig. S2.**
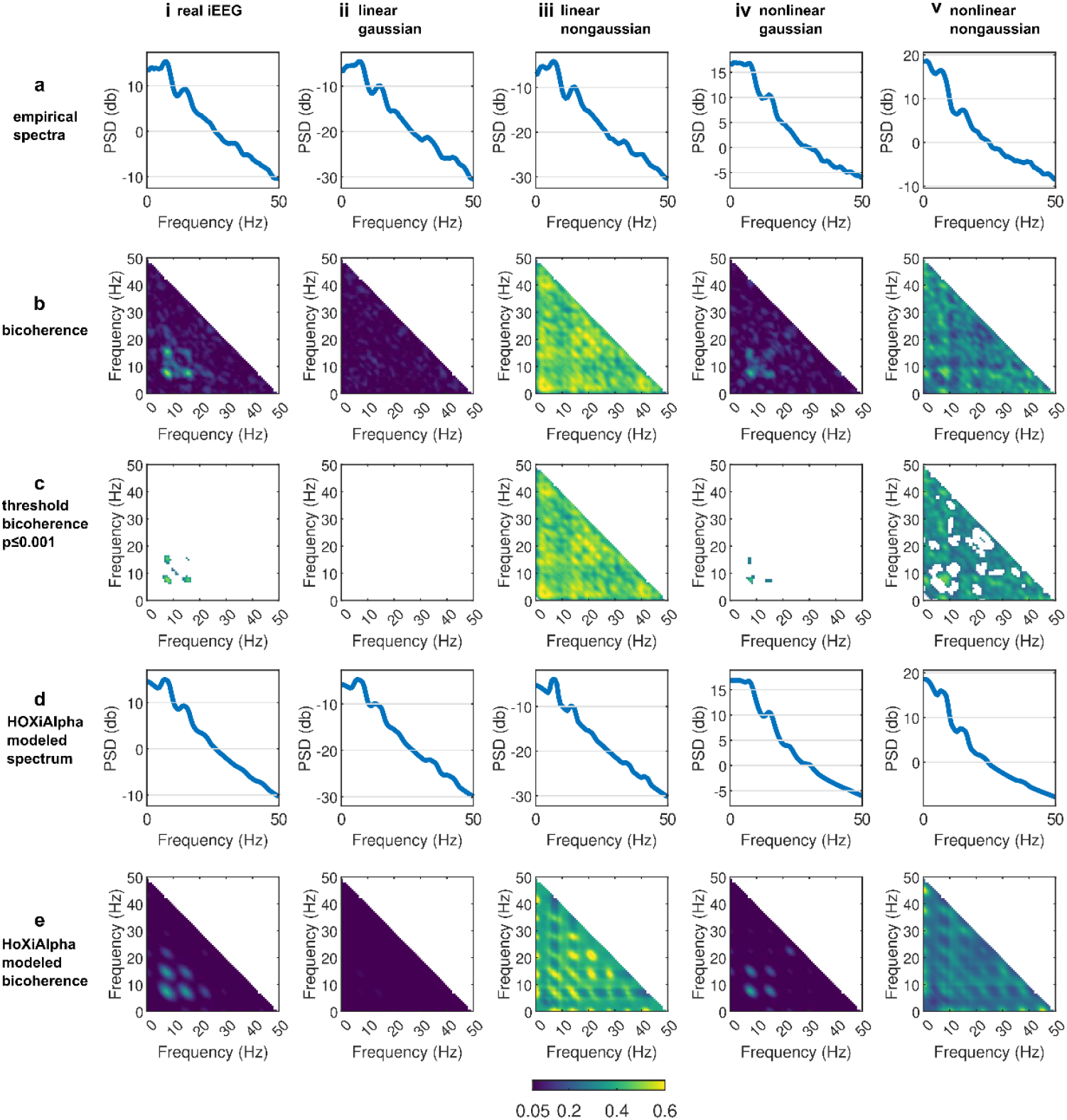
Simulation demonstration. Column from left to right are the 5 cases described in the simulation (i) real iEEG of MNI atlas data; (ii) linear Gaussian; (iii) linear non-Gaussian; (iv) nonlinear Gaussian; (v) nonlinear non-Gaussian; Row from top to bottom is the result of analysis of each step (a) empirical spectrum calculated with Multitaper method; b) empirical modulus bicoherence calculated with Multitaper method with color scale indicating bicoherence modulus amplitude.; (c) threshold bicoherence with parametric test of p≤0.001: providing a clearer view of which frequency interactions are nonlinearly coupled beyond test level; (d) the spectrum from the fitted BiSCA model; (e) the bicoherence from the fitted BiSCA model.

Fig. S 2 illustrates how non-Gaussianity and nonlinearity impact neural time series analyses; we synthesized intracranial EEG (iEEG) signals under five distinct scenarios—real iEEG data (MNI atlas), linear Gaussian, linear non-Gaussian, nonlinear Gaussian, and nonlinear non-Gaussian—and compared their power spectrum and bicoherence pattern. The real iEEG data Fig. S 2i exhibits a characteristic 1/f-like power spectrum with modest but significant bicoherence clusters after thresholding (p ≤ 0.001). This empirical baseline reveals that genuine neural signals naturally contain nonlinear and Gaussian input features.

In the Fig. S 3 we show the result of statistic test of Fig. S 2 Real iEEG (Top-Left): Both tests are significant, indicating the presence of both nonlinear interactions and non-Gaussian characteristics in the neural data. Linear Gaussian (Middle-Left): As expected for the null case, neither test is significant, correctly identifying the signal as linear and Gaussian. Linear Non-Gaussian (Middle-Right): Only the Gaussian test is significant, demonstrating the method’s ability to isolate non-Gaussian properties in the absence of nonlinearity. Nonlinear Gaussian (Bottom-Left): Only the linear test is significant, successfully detecting nonlinear dynamics driven by Gaussian innovations. Nonlinear Non-Gaussian (Bottom-Right): Both tests are significant, correctly identifying the joint presence of nonlinearity and non-Gaussianity. Overall, this figure demonstrates how the combined use of maximum and median bicoherence statistics provides a robust framework for disentangling the distinct contributions of nonlinearity and non-Gaussianity in complex time series.

**Fig. S3.**
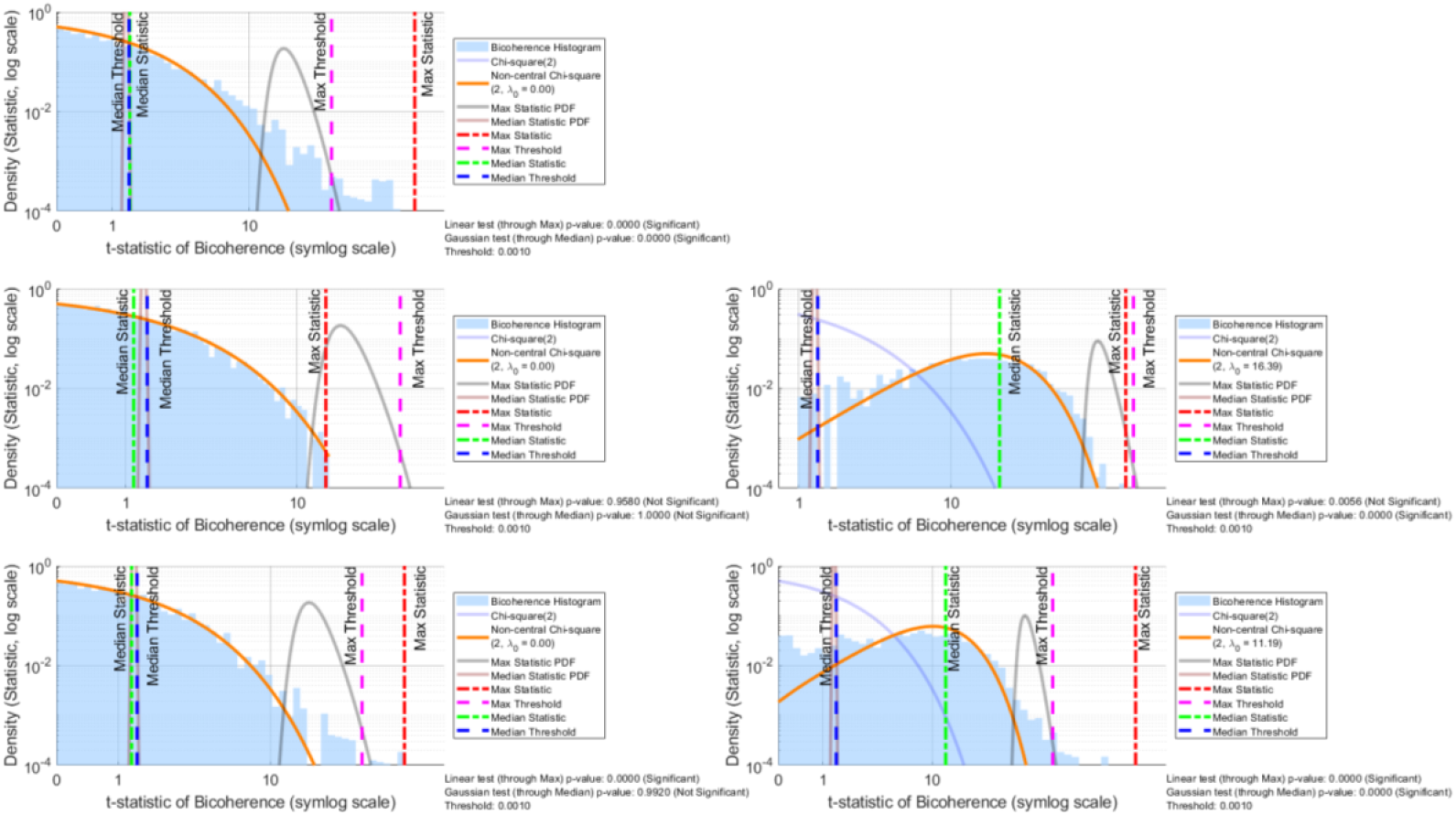
Statistical analysis of bicoherence for real and simulated time series. The figure displays the results of statistical tests applied to the bicoherence of five different time series signals to distinguish between nonlinearity and non-Gaussianity. The five cases are: (Top-left) Real intracranial EEG (iEEG) data; (Middle-left) A simulated linear Gaussian process; (Middle-right) A simulated linear non-Gaussian process; (Bottom-left) A simulated nonlinear Gaussian process; and (Bottom-right) A simulated nonlinear non-Gaussian process. Each subplot shows: Bicoherence Histogram (Light Blue): The distribution of the t-statistic calculated from the empirical bicoherence values of the signal. Chi-square Distributions: The theoretical central chi-square distribution with 2 degrees of freedom (blue line, *χ*^2^(2)), which is expected for a linear Gaussian process. A non-central chi-square distribution (orange line) is fitted to the data, where the non-centrality parameter (*λ*_0_) captures deviations from the null hypothesis, related to non-Gaussianity. Statistic PDFs (Gray and Dark Red): The probability density functions for the ‘Max’ and ‘Median’ statistics, representing the expected distribution under the null hypothesis. Vertical Lines: The observed values for the ‘Max Statistic’ (red) and ‘Median Statistic’ (green) are compared against their respective significance thresholds, ‘Max Threshold’ (magenta) and ‘Median Threshold’ (blue). The text at the bottom of each plot summarizes two hypothesis tests: 1. Linear test (through Max): Uses the maximum bicoherence value to test for significant quadratic phase coupling, which indicates nonlinearity. A significant result occurs if the ‘Max Statistic’ exceeds the ‘Max Threshold’. 2. Gaussian test (through median): Uses the median bicoherence value to test for a constant offset in bicoherence, which is characteristic of linear non-Gaussian processes. A significant result occurs if the ‘Median Statistic’ exceeds the ‘Median Threshold’.

The result shows that the bicoherence can be different even if the second-order spectrum is similar. In the linear non-Gaussian scenario Fig. S 2iii, the data deviate from Gaussianity but do not exhibit intrinsic nonlinear dependencies. This linear yet non-Gaussian structure typically manifests as a uniform or “constant” offset in the bicoherence^[74]^. In other words, rather than showing localized peaks at specific bifrequencies, the bicoherence tends to exhibit a more diffuse elevation across the entire frequency domain. Conversely, the bicoherence in the nonlinear Gaussian scenario is limited to well-defined regions (i.e., specific bifrequencies where nonlinearity introduces phase coupling), with otherwise negligible values. This difference underscores that nonlinearity and non-Gaussianity can contribute distinctly to the observed cross-frequency coupling patterns, helping to explain why the bicoherence in the linear non-Gaussian simulation diverges markedly from that of real iEEG data.

We further fitted each case with our BiSCA model, which accurately reproduced both the power spectral density and bicoherence for each scenario (rows of *d* and *e* in Fig. S 2). Notably, the model faithfully recapitulated the real iEEG features, mirroring the 1/f-like spectral decay and the subtle, significant cross-frequency interactions. These results highlight the model’s capacity to disentangle and reconstruct complex neural signals, shedding light on the interplay of Gaussianity and nonlinearity that shapes iEEG dynamics.

#### S.3.2 Simulation for the Geometric view of bicoherence

The simulation of Fig. 2 shows the geometric point of view of the bicoherence. The simulation employed an NNAR method to analyze nonlinear dynamics in intracranial iEEG data containing Wicket wave. The pipeline began with preprocessing a 2-second iEEG segment (sampled at 200 Hz) from a subject with normal variant activity. After reconstructing the phase space using a 23rd-order embedding, the NNAR model was trained to capture the system’s dynamics. A pruning step removed non-equilibrium states, retaining only key dynamics with weighted contributions exceeding 1e-2.

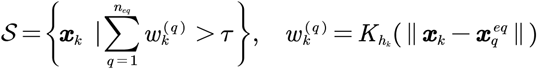

Deterministic and stochastic forecasts were generated using the pruned model. The deterministic (noise free) simulation to show the 3D phase portrait skeleton visualization, spectrum and bicoherence of this limit cycle. The stochastic forecasts with innovations modeled as Gaussian noise scaled by residual covariance to obtain potential landscape of 2D state space. The system’s geometry was projected into 2D/3D state spaces via PCA of the high dimensional states set 𝒮, and potential landscapes were estimated using kernel density methods on 100×100 grids, this is a nonparametric version of the landscape estimation^[101]^. Spectrum and bicoherence, was implemented also with Multitaper estimators (*NW* = 2.5) across 300-sample windows with 75% overlap.

To contrast with the asymmetric dynamics, we simulated symmetric nonlinear oscillations by enforcing the nonparametric state evolution function to satisfy *F* (−***x***) = − F(***x***),making the system dynamics invariant under state inversion (**X**⟶ − **X**). The modified state evolution becomes:

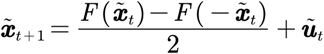

where the averaging ensures cancellation of even-order nonlinearities.

The reconstructed phase portrait revealed a stable limit cycle (Fig. 2C), confirming the system’s oscillatory nature. The potential landscape exhibited asymmetric minima (Fig. 2I), explaining the spikes’ directional morphology. Spectral analysis showed harmonically related peaks at 8.7Hz and 17.4Hz (Fig. 2J), with total harmonic distortion (THD) −4.08dB. Bicoherence analysis demonstrated significant phase coupling between fundamental and harmonic components (Fig. 2K), quantifying the nonlinear interactions. The symmetric version of the system, depicted in subplot (B) to (F), provides a contrasting perspective to the asymmetric case. Besides the symmetricity of the time domain analysis (B) to (D), the spectrum shows it only has the odd order harmonics, therefore, the bicoherence shows zero everywhere.

The asymmetric bicoherence profile (Fig. 2K) reflects directionally biased nonlinear interactions, characteristic of depolarization-hyperpolarization asymmetry in the neuronal networks. The reconstructed limit cycle (Fig. 2H) suggests possible self-sustaining oscillations stabilized by voltage-gated ion channel dynamics during the sample time interval. The potential landscape’s asymmetry (Fig. 2) aligns with the greater energy required for spike initiation versus termination, consistent with sodium channel inactivation kinetics. The 9 Hz spectral peak corresponds to thalamocortical resonance frequencies implicated in spike-wave generation. While the model successfully captured short-term dynamics (τ < 100 ms), longer simulations showed phase drift, this could be introduced by the step of pruning removed the activities shifted by the slow variables, the underlying attractor may not perfect limit cycle but quasiperiodic or chaotic, suggesting the need for additional slow variables in future extensions ^[102,103]^. These analysis bridges the spectral domain nonlinear analysis with the time domain evolution, providing a framework for quantifying stability landscapes in the nonlinear oscillations

The geometric interpretation of bicoherence provides critical insights into system dynamics. For an asymmetric system (Fig. 2G-K), significant bicoherence between the fundamental frequency (8.7 Hz) and its second harmonic (17.4 Hz) signifies quadratic phase coupling. This interlocking of phases constrains the system’s trajectory in phase space to a specific manifold, manifesting as an asymmetric “wicket” waveform and a distorted limit cycle in the phase portrait. Conversely, for a symmetric system (Fig. 2B-F), the bicoherence is zero, reflecting the absence of phase coupling. The system’s trajectory is unconstrained, resulting in a uniform limit cycle and a time-domain waveform with only odd-order harmonics. This contrast highlights how bicoherence geometrically quantifies the degree of asymmetry in the system’s underlying potential landscape. Furthermore, bicoherence helps distinguish system-based nonlinearity from non-Gaussian noise. In a nonlinear system, the energy landscape imposes state-dependent constraints, suppressing noise-induced perturbations more effectively in regions with strong recovery forces (e.g., steep potential wells). This state-specific “filtering” of non-Gaussian inputs reduces the measured bicoherence compared to a linear system, where such perturbations propagate homogeneously (Fig. 2). This distinction underscores the importance of modeling both system nonlinearity and noise non-Gaussianity when interpreting higher-order spectral features.

The geometric interpretation of bicoherence reveals critical insights into the nonlinear dynamics of the system, particularly in how phase coupling shapes its oscillatory behavior. In the asymmetric system, the bicoherence plot (Fig. 2K) shows significant phase coupling between the fundamental frequency at 8.7 Hz and its second harmonic at 17.4 Hz, indicating quadratic nonlinear interactions that constrain the system’s trajectory in phase space to a manifold reflecting these phase-locked oscillations. This interlocking of phases manifests as the asymmetric morphology in the time-domain waveform of the Wicket (Fig. 2A), with the directional bias corresponding to imbalanced phase relationships. The reconstructed phase portrait Fig. 2H) supports this, as these nonlinear constraints shape the stable limit cycle’s trajectory. Conversely, in the symmetric system, the bicoherence is zero everywhere (Fig. 2F), aligning with the regular, balanced oscillations in the time domain (Fig. 2B) and the presence of only odd-order harmonics (Fig. 2E). Here, the absence of phase coupling results in an unconstrained trajectory in phase space, producing a more uniform limit cycle (Fig. 2C). This contrast highlights how bicoherence geometrically quantifies the degree of nonlinearity and asymmetry in the system’s dynamics, linking spectral domain insights to the time-domain evolution and the underlying potential landscape (Fig. 2D and Fig. 2I). These findings underscore the utility of bicoherence in analyzing stability and morphology in nonlinear oscillatory systems, offering a framework to explore the interplay between structure and dynamics.

### S.4 Goodness of fit

To demonstrate how much variance can be explained by the model, here we put the cumulative density function of the ***R***^2^. In the formula defined in formula (13) and (15), the parameter {*μ*_*ρk*_} has two options: 1) Harmonic fixed with the harmonic relationship of the “fundamental oscillation”, 2) Free: initialize with spectrum and optimize the joint fit. We put the ***R***^2^ of both cases here.

**Fig. S4.**
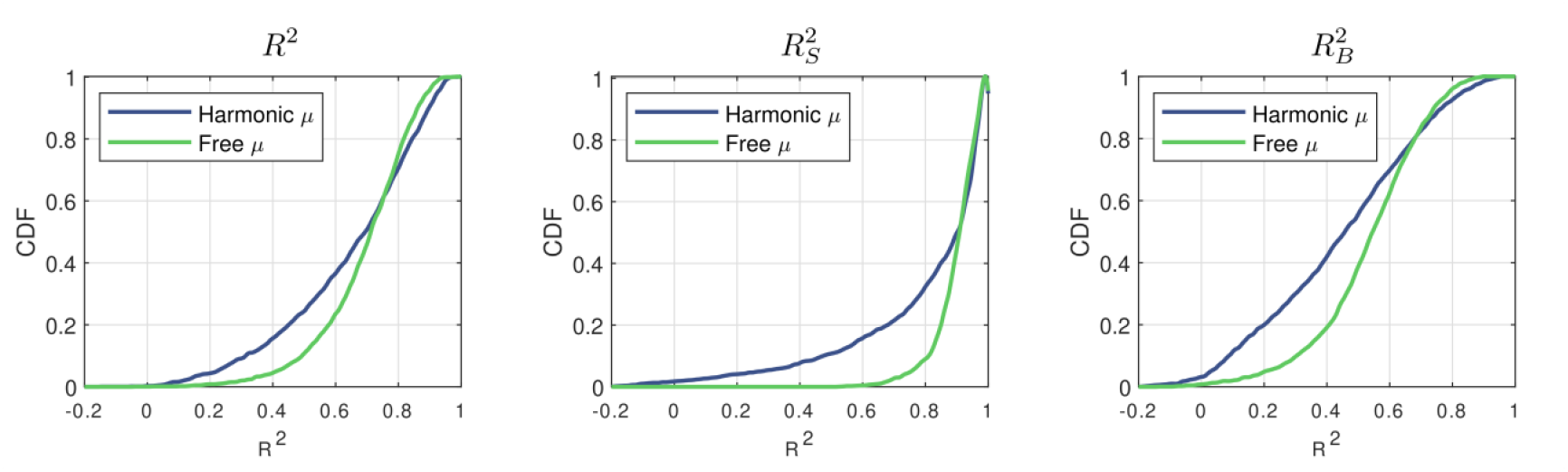
Distributions of the coefficient of determination R^2^ in three different conditions. a) is the **R**^2^ of the full model; b) is the **R**^2^ for the spectrum; c) is the **R**^2^ for the bispectrum

The ***R***^2^ distribution is calculated for the fit of each channel shown in the Fig. S 4. The goodness of fit suggests that the model’s goodness of fit varies considerably across the entire dataset. The “free” option of the peak center fits with a greater degree of freedom. Therefore, its ***R***^2^ is concentrated on the right side, compared to the “harmonic” organized center peak option. Some phenomena, such as split Alpha, may not be described by the “harmonic” organized model.

Fig. S 4A shows a broad distribution of overall ***R***^2^ values, suggesting that the model’s goodness of fit varies considerably across the entire dataset. A noticeable upward trend toward higher ***R***^2^ (up to ≈) indicates that many data segments exhibit strong fits, although a small portion remains at the lower end of the scale. Since the variance of the spectrum and the variance of the bispectrum is quite different, which also vary with subject the 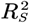 and 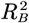 are more intuitive. Fig. S 4B 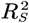 are comparatively narrower near the high value but rise sharply toward 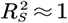. This pattern implies that many single-subject spectral fits are quite robust, with fewer cases in the middle range of goodness of fit. Fig. S 4C presents a more bell-shaped distribution of baseline 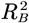 values. Most of these are clustered around moderate goodness-of-fit levels 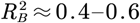, with fewer instances at the extremes (near 0 or near 1). Overall, bispectrum fits appear to be consistent but rarely reach near-perfect levels.

### S.5 Peak relation from the spectrum

**Fig. S5.**
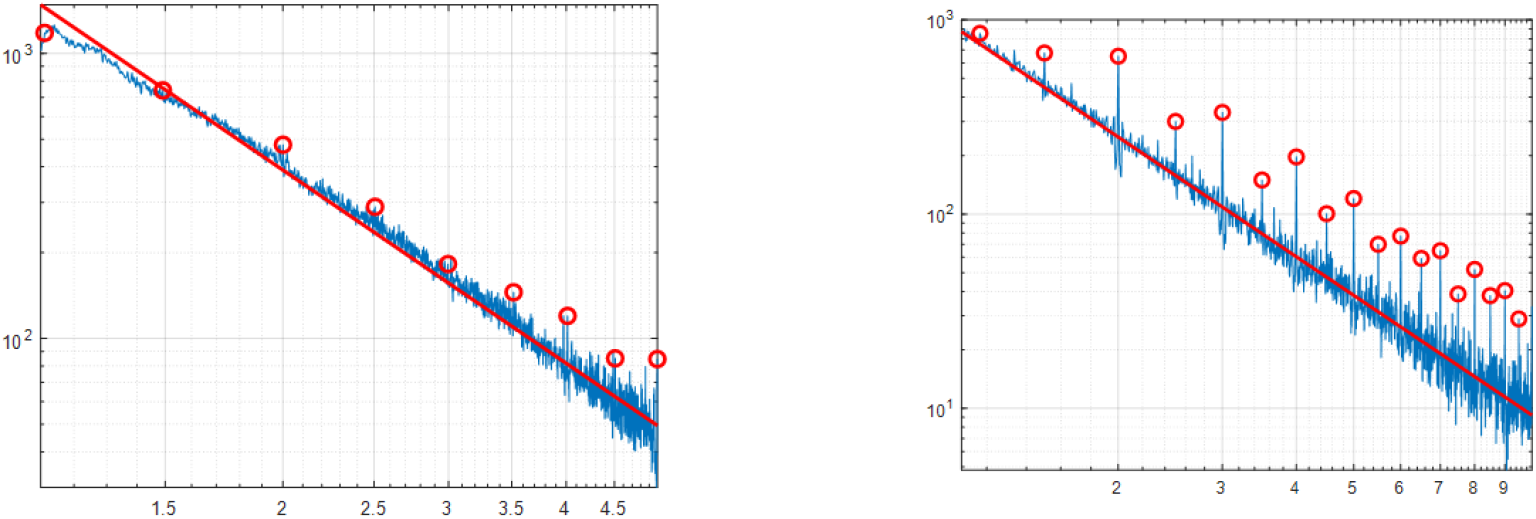
the ratio of the peaks identified in the spectrum. The left is from harmonized 1563 × 18 channels scalp EEG data. The right is from 1771 channels of iEEG data.

Besides integer harmonic peaks of periodic oscillations, other proportions between peak frequency reveal possible quasi-periodic motion and chaotic; the distribution of the proportions showed possible scale free of the organization of peaks. On the Exponent Scale, the slope is 2.2531 for EEG and 2.0493 for iEEG. However, metaphysical models attempt to impose intriguing physical interpretations on empirical data without necessarily ensuring that these interpretations adhere to rigorous statistical or biophysical validity. Evertz et al. (2022)^[104]^ point out correctly that there is doubt that the aperiodic component is either “scale-free” or a reflection of criticality. An alternative is to look at biophysical models with effects of the extracellular space ^[105]^ as mentioned by Evertz et al. (2022)^[104]^. In addition to interest and questions about the scale-free nature of ξ process ^[106]^. This question was discussed in Valdés-Sosa et al. in 1999^[100,107]^, when people argue that the EEG is chaos, the neural activity is a nonlinear stochastic system driven by the noise. We reserve these possibilities, and these conclusions still need to be verified by rigorous statistical methods. One possibility is that using the bicoherence in this paper to study the relationship between the components is possible when all the ratios here are the intermodulation combination of the integer peaks.

## References

[1] E. Nozari, M. A. Bertolero, J. Stiso, L. Caciagli, E. J. Cornblath, X. He, A. S. Mahadevan, G. J. Pappas, D. S. Bassett, Nat. Biomed. Eng. 2024, 8, 68.

[2] G. T. Raffaelli, S. Jiøíèek, J. Hlinka, Nonlinear brain connectivity from neurons to networks: quantification, sources and localization, bioRxiv 2024.

[3] M. Breakspear, Nat. Neurosci. 2017, 20, 340.

[4] C. J. Stam, J. P. M. Pijn, P. Suffczynski, F. H. Lopes da Silva, Clinical Neurophysiology 1999, 110, 1801.

[5] M. Breakspear, J. R. Terry, Clinical Neurophysiology 2002, 113, 735.

[6] S. R. Cole, B. Voytek, Trends Cogn Sci 2017, 21, 137.

[7] F. H. Lopes da Silva, A. van Rotterdam, P. Barts, E. van Heusden, W. Burr, in Progress in Brain Research (Eds.: M. A. Corner, D. F. Swaab), Vol. 45, Elsevier 1976, pp. 281–308.

[8] L. H. Zetterberg, L. Kristiansson, K. Mossberg, Biol. Cybernetics 1978, 31, 15.

[9] F. Grimbert, O. Faugeras, Neural Computation 2006, 18, 3052.

[10] A. Spiegler, S. J. Kiebel, F. M. Atay, T. R. Knösche, NeuroImage 2010, 52, 1041.

[11] A. Bender, B. Voytek, N. Schaworonkow, Biorxiv 2025, 2023.10.13.562301.

[12] S. R. Cole, R. van der Meij, E. J. Peterson, C. de Hemptinne, P. A. Starr, B. Voytek, J. Neurosci.: Off. J. Soc. Neurosci. 2017, 37, 4830.

[13] S. Cole, B. Voytek, J. Neurophysiol. 2019.

[14] H. Carlqvist, V. V. Nikulin, J. O. Strömberg, T. Brismar, Med. Biol. Eng. Comput. 2005, 43, 599.

[15] T. Donoghue, N. Schaworonkow, B. Voytek, Eur. J. Neurosci. 2022, 55, 3502.

[16] M. J. Idaji, J. Zhang, T. Stephani, G. Nolte, K.-R. Müller, A. Villringer, V. V. Nikulin, NeuroImage 2022, 252, 119053.

[17] W. Klimesch, Trends in Cognitive Sciences 2012, 16, 606.

[18] M. Labecki, R. Kus, A. Brzozowska, T. Stacewicz, B. S. Bhattacharya, P. Suffczynski, Frontiers in Computational Neuroscience 2016, 10.

[19] K. A. Mohsini, O. Farooq, Y. U. Khan, M. Tripathi, in 2017 International Conference on Multimedia, Signal Processing and Communication Technologies (IMPACT), 2017, pp. 209–213.

[20] J. A. Roberts, P. A. Robinson, NeuroImage 2012, 62, 1947.

[21] N. Schaworonkow, Imaging Neuroscience 2023, 1, 1.

[22] S. Van Albada, P. Robinson, Frontiers in Human Neuroscience 2013, 7.

[23] S. Bartz, F. S. Avarvand, G. Leicht, G. Nolte, NeuroImage 2019, 188, 145.

[24] C. S. Zandvoort, G. Nolte, Journal of Neuroscience Methods 2021, 350, 109032.

[25] V. Jirsa, V. Müller, Front. Comput. Neurosci. 2013, 7.

[26] H. T. Zhu, ASCE-ASME Journal of Risk and Uncertainty in Engineering Systems, Part B: Mechanical Engineering 2015, 1, 011005.

[27] D. R. Brillinger, Time Series: Data Analysis and Theory, SIAM: Society for Industrial and Applied Mathematics 2001.

[28] R. D. Pascual-marqui, P. A. Valdes-sosa, A. Alvarez-amador, International Journal of Neuroscience 1987, 40, 89.

[29] T. Donoghue, M. Haller, E. J. Peterson, P. Varma, P. Sebastian, R. Gao, T. Noto, A. H. Lara, J. D. Wallis, R. T. Knight, A. Shestyuk, B. Voytek, Nature Neuroscience 2020, 23, 1655.

[30] C. W. Connor, Anesth. Analg. 2022, 10.1213/ANE.7577.

[31] S. Z. Dragovic, J. Ostertag, N. Baumann, P. S. García, S. Kratzer, G. Schneider, S. Schwerin, J. Sleigh, M. Kreuzer, Anesth. Analg. 2022, 10.1213/ANE.7530.

[32] S. D. McKeon, M. I. Perica, A. C. Parr, F. J. Calabro, W. Foran, H. Hetherington, C.-H. Moon, B. Luna, Dev. Cognit. Neurosci. 2024, 66, 101373.

[33] N. Brake, F. Duc, A. Rokos, F. Arseneau, S. Shahiri, A. Khadra, G. Plourde, Nat Commun 2024, 15, 1514.

[34] M. A. Kramer, C. J. Chu, Neural Comput. 2024, 36, 1643.

[35] L. V. Marcuse, M. C. Fields, J. J. Yoo, Rowan’s Primer of EEG, 2nd edition., Elsevier, Edinburgh 2015.

[36] M. J. Hinich, Journal of Time Series Analysis 1982, 3, 169.

[37] Y. Wang, M. Li, D. Paz-Linares, A. Areces-Gonzalez, M. Bringas-Vega, J. Bosch-Bayard, P. Valdes-Sosa, in The Organization for Human Brain Mapping (OHBM) 2024 Annual Meeting, 2024, p. 1657.

[38] N. Schaworonkow, V. V. Nikulin, PLoS Comput Biol 2019, 15, e1007055.

[39] N. Schaworonkow, V. V. Nikulin, NeuroImage 2022, 253, 119093.

[40] R. D. Pascual-Marqui, K. Kochi, T. Kinoshita, Cortical Xi-Alpha model for resting state electric neuronal activity, arXiv 2022.

[41] L. R. Silva, Y. Amitai, B. W. Connors, Science 1991, 251, 432.

[42] T. Van Kerkoerle, M. W. Self, B. Dagnino, M.-A. Gariel-Mathis, J. Poort, C. Van Der Togt, P. R. Roelfsema, Proc. Natl. Acad. Sci. 2014, 111, 14332.

[43] D. Mendoza-Halliday, A. J. Major, N. Lee, M. J. Lichtenfeld, B. Carlson, B. Mitchell, P. D. Meng, Y. (Sophy) Xiong, J. A. Westerberg, X. Jia, K. D. Johnston, J. Selvanayagam, S. Everling, A. Maier, R. Desimone, E. K. Miller, A. M. Bastos, Nat. Neurosci. 2024, 27, 547.

[44] N. Brake, F. Duc, A. Rokos, F. Arseneau, S. Shahiri, A. Khadra, G. Plourde, Nat Commun 2024, 15.

[45] P. F. Bloniasz, S. Oyama, E. P. Stephen, Filtered point processes tractably capture rhythmic and broadband power spectral structure in neural electrophysiological recordings, Cold Spring Harbor Laboratory 2024.

[46] P. Marzocca, J. M. Nichols, A. Milanese, M. Seaver, S. Trickey, Mechanical Systems and Signal Processing 2008, 22, 1882.

[47] J. M. Nichols, C. C. Olson, J. V. Michalowicz, F. Bucholtz, IEEE Transactions on Signal Processing 2009, 57, 3879.

[48] A. A. Amador, R. D. Pascual-Marqui, P.A. Valdés-Sosa, in Machinery of the Mind: Data, Theory, and Speculations About Higher Brain Function (Eds.: E. R. John, T. Harmony, L. S. Prichep, M. Valdés-Sosa, P.A. Valdés-Sosa), Birkhäuser, Boston, MA 1990, pp. 59–90.

[49] P. Valdés, J. Bosch, R. Grave, J. Hernandez, J. Riera, R. Pascual, R. Biscay, Brain Topogr 1992, 4, 309.

[50] F. Freyer, J. A. Roberts, R. Becker, P. A. Robinson, P. Ritter, M. Breakspear, J. Neurosci. 2011, 31, 6353.

[51] F. Freyer, K. Aquino, P. A. Robinson, P. Ritter, M. Breakspear, J. Neurosci. 2009, 29, 8512.

[52] S. H. Wang, F. Siebenhühner, G. Arnulfo, V. Myrov, L. Nobili, M. Breakspear, S. Palva, J. M. Palva, J. Neurosci. 2023, 43, 7642.

[53] B. Frauscher, N. von Ellenrieder, R. Zelmann, I. Doležalová, L. Minotti, A. Olivier, J. Hall, D. Hoffmann, D. K. Nguyen, P. Kahane, F. Dubeau, J. Gotman, Brain 2018, 141, 1130.

[54] F. Shahbazi Avarvand, S. Bartz, C. Andreou, W. Samek, G. Leicht, C. Mulert, A. K. Engel, G. Nolte, NeuroImage 2018, 174, 352.

[55] J. Pfanzagl, Metrika 1969, 14, 249.

[56] J. Rice, Journal of Multivariate Analysis 1979, 9, 378.

[57] N. N. Leonenko, A. Y. Sikorskii, G. Terdik, 1998, 6, 159.

[58] V. V. Anh, N. N. Leonenko, L. M. Sakhno, Journal of Statistical Planning and Inference 2004, 123, 161.

[59] V. V. Anh, N. N. Leonenko, L. M. Sakhno, Journal of Statistical Planning and Inference 2007, 137, 1302.

[60] V. V. Anh, N. N. Leonenko, L. M. Sakhno, Journal of Multivariate Analysis 2007, 98, 706.

[61] L. Sakhno, in Modern Stochastics and Applications (Eds.: V. Korolyuk, N. Limnios, Y. Mishura, L. Sakhno, G. Shevchenko), Springer International Publishing, Cham 2014, pp. 319–336.

[62] L. Sakhno, Austrian Journal of Statistics 2020, 49, 106.

[63] F. Freyer, J. A. Roberts, P. Ritter, M. Breakspear, PLoS Comput Biol 2012, 8, e1002634.

[64] Y. Wang, M. Li, R. G. Reyes, Ronaldo García Reyes, J. Bosch-Bayard, M. Bringas-Vega, Michael Breakspear, L. Minati, P. Valdes-Sosa, in The Organization for Human Brain Mapping (OHBM) 2025 Annual Meeting, 2025, p. 1339.

[65] M. Li, Y. Wang, C. Lopez-Naranjo, R. C. G. Reyes, A. I. A. Hamid, A. C. Evans, A. N. Savostyanov, A. Calzada-Reyes, A. Areces-Gonzalez, A. Villringer, C. A. Tobon-Quintero, D. Garcia-Agustin, D. Paz-Linares, D. Yao, L. Dong, E. Aubert-Vazquez, F. Reza, H. Omar, J. M. Abdullah, J. R. Galler, J. F. Ochoa-Gomez, L. S. Prichep, L. Galan-Garcia, L. Morales-Chacon, M. J. Valdes-Sosa, M. Tröndle, M. F. B. M. Zulkifly, M. R. B. A. Rahman, N. S. Milakhina, N. Langer, P. Rudych, S. Hu, T. Koenig, T. A. Virues-Alba, X. Lei, M. L. Bringas-Vega, J. F. Bosch-Bayard, P. A. Valdes-Sosa, NeuroImage 2022, 119190.

[66] D. J. Thomson, Proceedings of the IEEE 1982, 70, 1055.

[67] H. He, D. J. Thomson, J Comput Neurosci 2010, 29, 23.

[68] K. S. Riedel, A. Sidorenko, IEEE Transactions on Signal Processing 1995, 43, 188.

[69] D. R. Brillinger, M. Rosenblatt, Proceedings of the National Academy of Sciences 1967, 57, 206.

[70] D. R. Brillinger, M. Rosenblatt, 1967.

[71] D. J. Thomson, in Workshop on Higher-Order Spectral Analysis, 1989, pp. 19–23.

[72] Y. Birkelund, A. Hanssen, E. J. Powers, in 2001 IEEE International Conference on Acoustics, Speech, and Signal Processing. Proceedings (Cat. No.01CH37221), Vol. 5, 2001, pp. 3085–3088 vol.5.

[73] F. Shahbazi, A. Ewald, G. Nolte, J Neurosci Methods 2014, 233, 177.

[74] C. Nikias, A. Petropulu, Higher Order Spectra Analysis: A Non-Linear Signal Processing Framework, Facsimile edition., Pearson, Englewood Cliffs, N.J 1993.

[75] R. D. Pascual-marqui, P. A. Valdes-sosa, A. Alvarez-amador, International Journal of Neuroscience 1988, 40, 89.

[76] M. Newville, T. Stensitzki, D. B. Allen, A. Ingargiola, LMFIT: Non-Linear Least-Square Minimization and Curve-Fitting for Python, Zenodo 2014.

[77] C. L. Haley, M. Anitescu, IEEE Signal Process Lett. 2017, 24, 1696.

[78] S. A. Billings, Nonlinear System Identification: NARMAX Methods in the Time, Frequency, and Spatio-Temporal Domains, 1st ed., Wiley 2013.

[79] Z. K. Peng, W. M. Zhang, B. T. Yang, G. Meng, F. L. Chu, Mechanical Systems and Signal Processing 2013, 36, 456.

[80] X. Jing, Z. Lang, Frequency Domain Analysis and Design of Nonlinear Systems based on Volterra Series Expansion: A Parametric Characteristic Approach, Springer International Publishing, Cham 2015.

[81] C. L. Nikias, J. M. Mendel, IEEE Signal Processing Magazine 1993, 10, 10.

[82] Bevan S., Kullberg R., Rice J., The Annals of Statistics 1979, 7, 237.

[83] D. R. Brillinger, J Am Water Resources Assoc 1985, 21, 743.

[84] A. Travert, C. Fernandez, SpectroChemPy, Zenodo 2024.

[85] H. E. Lackey, R. L. Sell, G. L. Nelson, T. A. Bryan, A. M. Lines, S. A. Bryan, J. Chem. Educ. 2023, 100, 2608.

[86] A. M. Price-Whelan, P. L. Lim, N. Earl, Others, Astrophys, J. 2022, 935, 167.

[87] D. A. Guimarães, in Anais do XXXIX Simpósio Brasileiro de Telecomunicações e Processamento de Sinais, Sociedade Brasileira de Telecomunicações 2021.

[88] L. H. Zetterberg, Mathematical Biosciences 1969, 5, 227.

[89] A. Isaksson, K. Lagergren, A. Wennberg, Electroencephalography and Clinical Neurophysiology 1976, 41, 225.

[90] S. V. Narasimhan, Signal Process. 1989, 18, 17.

[91] A. M. Hughes, T. A. Whitten, J. B. Caplan, C. T. Dickson, Hippocampus 2012, 22, 1417.

[92] J. Q. Kosciessa, T. H. Grandy, D. D. Garrett, M. Werkle-Bergner, NeuroImage 2020, 206, 116331.

[93] R. A. Seymour, N. Alexander, E. A. Maguire, European Journal of Neuroscience 2022, 56, 5836.

[94] H. Wen, Z. Liu, Brain Topogr 2016, 29, 13.

[95] L. E. Wilson, J. da Silva Castanheira, S. Baillet, eLife 2022, 11, e77348.

[96] R. J. Barry, F. M. D. Blasio, J. Neural Eng. 2021, 18, 034001.

[97] S. Hu, Z. Zhang, X. Zhang, X. Wu, P. A. Valdes-Sosa, IEEE J. Biomed. Health Inform. 2024, 28, 2624.

[98] R. G. Reyes, A. A. Gonzales, Y. Wang, Y. Jin, L. Minatti, P. A. Valdes-Sosa, Lifespan mapping of EEG source spectral dynamics with ξ − αNET, bioRxiv 2025.

[99] J. R. M. Hosking, J. R. Wallis, Regional Frequency Analysis: An Approach Based on L-Moments, Cambridge University Press, Cambridge 1997.

[100] P. Valdés-Sosa, J. Bosch, J. Jiménez, N. Trujillo-Barreto, R. J. Lirio, F. Morales, J. luis Hernandez Caceres, T. Ozaki, in Non linear Dynamic and Brain Funcioning, 1999, pp. 278–284.

[101] R. Zhou, Y. Yu, C. Li, Iscience 2024, 27.

[102] K. M. Stiefel, G. B. Ermentrout, J. Neurophysiol. 2016, 116, 2950.

[103] M. Breakspear, Nat. Neurosci. 2017, 20, 340.

[104] R. Evertz, D. G. Hicks, D. T. J. Liley, PLOS Computational Biology 2022, 18, e1010012.

[105] H. Lindén, K. H. Pettersen, G. T. Einevoll, J Comput Neurosci 2010, 29, 423.

[106] C. Bédard, H. Kröger, A. Destexhe, Phys. Rev. Lett. 2006, 97, 118102.

[107] P. A. Valdes, J. C. Jimenez, J. Riera, R. Biscay, T. Ozaki, Biological Cybernetics 1999, 81, 415.

